# Benchmarking Short-Read ITS2 and Full-Length ITS Sequencing Reveals Pipeline-Dependent Biases in Indoor Fungal Community Profiling

**DOI:** 10.64898/2026.05.15.725464

**Authors:** Mengyi Dong, Denene Blackwood, Megan Lott, Sherlynette Pérez Castro, Xavier Larkin, Thomas J. Clerkin, Heather Hemric, Jake Nash, Yeon Ji Kim, Jason W. Arnold, Lawrence A. David, Rytas J. Vilgalys, Anthony A. Fodor, Rachel T. Noble

## Abstract

Short-read amplicon sequencing is widely used for fungal surveys but can limit taxonomic resolution. Long-read sequencing enables recovery of the full internal transcribed spacer (ITS) region and may improve ecological and taxonomic inference. Here, we conducted a paired comparison of Illumina ITS2 and PacBio HiFi full-length ITS sequencing using identical DNA extracts from built-environmental air and surface samples (n = 68) collected across homes, a dormitory, and laboratories. Both datasets were taxonomically assigned using the same algorithm and reference database. We performed paired statistics, in-silico ITS2 trimming of long-read sequences, and cross-platform mapping at multiple identity thresholds. Full-length ITS provided higher taxonomic resolution, assigning a greater fraction of ASVs at the family (98% vs. 88%) and species (42% vs. 32%) ranks than ITS2 (paired Wilcoxon q = 0.002). Alpha-diversity comparisons showed similar Shannon diversity across pipelines, whereas richness metrics were consistently higher for full-length ITS. Beta-diversity analyses indicated broadly comparable community-level patterns, although full-length ITS revealed stronger sample-type- and location-associated structure (PERMANOVA R² ≥ 0.06, p = 0.0001). In-silico ITS2 trimming reduced these differences, indicating that amplicon length is a major contributor to enhanced taxonomic resolution and ecological inference. Cross-platform mapping further showed extensive one-to-many relationships between ITS2 and full-length ITS ASVs, consistent with increased sequence resolution in long-read data.

Together, these results show that ITS2 sequencing provides robust community-level profiling, while full-length ITS enables improved richness estimates and finer ecological and taxonomic resolution. This paired, bias-aware framework provides a practical template for selecting fungal amplicon sequencing strategies in built-environment mycobiome studies.

**Importance:** Fungal communities in built environments influence indoor air quality and human exposure, yet their characterization depends strongly on sequencing strategy. This study provides a controlled, paired comparison of short-read ITS2 and long-read full-length ITS sequencing, showing that differences in amplicon length substantially contribute to variation in taxonomic resolution and ecological inference. While both approaches yield comparable community-level patterns, full-length ITS improves richness estimates, species-level assignment, and environmental discrimination by resolving sequence variation collapsed in ITS2 surveys. By integrating paired diversity analyses, in-silico ITS2 trimming, and cross-platform ASV mapping, this work offers a bias-aware framework for evaluating fungal amplicon pipelines. Importantly, improved species-level resolution enables functional interpretation of indoor fungi, for example the identification of taxa associated with pathogenic traits, allergen production, or toxin synthesis, supporting the development of more informative exposure metrics and targeted assays relevant to human health in built environments.

## Introduction

The built-environment microbiome influences indoor air quality, building integrity, and human health (1). While indoor bacterial communities have been extensively characterized (1, 2), growing attention has focused on fungal diversity, spatial variation, and exposure pathways in indoor environments (3–5). These studies show that indoor fungi originate from multiple sources and vary widely across buildings, yet accurate detection and classification of indoor fungal taxa remain challenging (1, 6).

A key challenge is the reliable identification of fungi at finer taxonomic ranks, especially for genera and species relevant to health. Indoor fungi span common saprophytes and opportunistic pathogens, with detection influenced by sampling approach, primer design, and sequencing platform (6, 7). Importantly, clinically relevant differences in pathogenicity and drug susceptibility are often observed among closely related fungal species or strains that are indistinguishable using coarse taxonomic markers. Well-documented examples include cryptic *Aspergillus* species within the *A. fumigatus* complex and members of the *Cryptococcus neoformans*/*C. gattii* species complexes (8). Indoor fungal exposure has been linked to respiratory health effects, particularly in children (9). Sequencing surveys routinely detect hundreds of fungal taxa within single homes, including medically important species such as *Aspergillus fumigatus*, *Stachybotrys chartarum*, and *Cryptococcus neoformans* (10–14). Sensitive and accurate molecular tools are therefore needed to interpret exposure risks and support environmental assessment (9).

Indoor fungal community composition varies across building types, ventilation, moisture, and occupant activity (15, 16). These factors shape both dominant indoor genera and lower-abundance taxa of potential concern. Such variability highlights the need for reproducible and comparable indoor fungal datasets (7).

Most environmental molecular fungal surveys rely on amplification of the internal transcribed spacer (ITS) region, with Illumina ITS1 or ITS2 sequencing widely used due to cost and throughput (17). ITS2 sequencing supports broad community profiling but often limits species-level resolution for closely related taxa (18). Long-read high-fidelity platforms such as PacBio HiFi can generate full-length ITS reads that retain additional phylogenetic information (19). Previous studies report improved taxonomic resolution with long-read ITS sequencing, alongside challenges related to primer design and performance across taxa (19–21). Recent improvements to PacBio sequencing throughput coupled with lower costs have made full-length ITS profiling more feasible for larger studies (22).

Despite the availability of both short- and long-read approaches, most indoor mycobiome studies rely on a single amplicon region (ITS1 or ITS2) (3, 5, 7). Few paired studies directly compare ITS2 and full-length ITS using the same indoor samples, and the relative contributions of amplicon length, primer choice, sequencing platform, and taxonomic classification remain unclear.

Here, we perform a paired comparison of Illumina ITS2 and PacBio HiFi full-length ITS sequencing to evaluate method-dependent differences in indoor fungal community profiling. Specifically, we compared taxonomic assignment across ranks under classifier parity, assessed agreement in community structure and genus-level abundance, evaluated patterns among health-relevant taxa, and used in-silico ITS2 trimming and cross-platform mapping to isolate the influence of amplicon length and primer targeting. Together, this work provides a method-focused benchmark for selecting fungal amplicon strategies in built-environment studies.

## Materials and Methods

### Sample collection and preprocessing

Indoor air and surface samples (n = 68) were collected between 2023 and 2024 from multiple built environments in North Carolina, including two unoccupied residences, a student dormitory, and laboratory spaces at the University of North Carolina at Chapel Hill and Duke University (Supplementary Data 1, Fig. S1). Sampling locations represented a range of occupancy levels, moisture conditions, and visible fungal growth. Surface swabs and bioaerosol samples were collected using standardized protocols appropriate to each site.

Surface samples were collected using sterile rayon swabs pre-moistened with buffer and swabbed over standardized surface areas. Bioaerosol samples were collected using a BobCat (AC-200) sampler with dry electret filters and eluted following manufacturer recommendations. Field blanks, method blanks, negative controls, and positive controls were included at each site to monitor contamination and processing consistency. All eluates were aliquoted and stored at −80 °C prior to DNA extraction. Detailed sampling locations, surface types, and control descriptions are provided in Supplementary Methods.

### DNA extraction and quality control

Total nucleic acids were extracted using magnetic-bead-based extraction on a KingFisher™ Flex system (bioMérieux NucliSENS kit). Extractions were eluted in 100 µL of elution buffer. Negative extraction controls and positive controls spiked with *Aspergillus niger* were included to assess contamination and extraction efficiency. All extracts were stored at −80 °C until library preparation. Extraction scripts and additional quality-control details are provided in the Supplementary Methods.

### Illumina ITS2 Library Preparation and Sequencing

The fungal ITS2 region was amplified using a three-step PCR protocol adapted from prior work (23) with ITS3NGS and ITS4NGR primers (24–26). All primer sequences are listed in S. Table 2. Amplicon sizes (∼400 bp) were verified by agarose gel electrophoresis and purified using AMPure XP Beads (Beckman Coulter, Inc., Brea, CA, USA) at a 1.8× bead-to-sample volume ratio. DNA concentrations were quantified using the Qubit HS dsDNA kit, and samples were pooled in equimolar amounts to ∼2–3 ng/µL. Final libraries were sequenced on an Illumina MiniSeq platform (Illumina, Inc., San Diego, CA, USA) targeting a read depth of at least 20,000 reads per sample using 2×250 bp paired-end reads in the Duke Sequencing and Genomic Technologies Shared Resource (Duke University, Durham, NC, USA). Detailed procedure and PCR cycling conditions are described in Supplementary Methods.

### PacBio Full-Length ITS Library Preparation and Sequencing

Full-length ITS amplicons were generated from the same DNA extracts using Phusion Plus polymerase with ITS1catta and ITS4ngsUni primers (20, 27). Samples were barcoded, pooled, and prepared following the PacBio Kinnex protocol without modification (28). Size selected and cleaned libraries were loaded onto a PacBio SMRT® Cell and sequenced on the Revio system (Pacific Biosciences of California, Inc.) at the Duke Sequencing and Genomic Technologies Shared Resource. Amplicon verification, pooling volumes, and circularization steps are described in the Supplementary Methods.

### Illumina Data Processing

Illumina paired-end reads were merged using PEAR (29), and ITS2 regions were extracted using ITSxpress (30). Sequences were processed in QIIME 2 (v2023.9) (31) using DADA2 denoise-single for error correction, chimera removal, and ASV inference (32). Taxonomy was assigned using a naïve Bayes classifier trained on full-length ITS in UNITE v9.0 (33). QIIME 2 artifacts were exported for downstream analysis in R.

### PacBio Data Processing

PacBio circular consensus sequences were processed using DADA2 v1.28 with PacBio-specific error modeling (32). Primers were removed prior to filtering, and ASVs were inferred with chimera removal optimized for long amplicons. Taxonomy was assigned using the same UNITE v9.0 classifier applied to the ITS2 data, ensuring classifier parity.

### Mapping of Illumina ITS2 ASVs Against PacBio Full-Length ITS References

Illumina ITS2 ASVs were mapped to PacBio full-length ITS ASVs using the official NCBI BLAST + Docker image (v2.11.0; https://hub.docker.com/r/ncbi/blast) at >97% identity to assess sequence overlap and resolution differences. Detailed parameters are provided in the Supplementary Methods. Maximum likelihood trees were constructed for the BLAST mapped sequences. For downstream mapping analysis, identity thresholds of ≥99% or 100% were used to quantify one-to-one and one-to-many relationships between pipelines.

### In-silico ITS2 Extraction from PacBio Full-Length ITS Reads

An in-silico ITS2 dataset was generated from PacBio full-length ITS ASVs using ITSx (34). Extracted ITS2 sequences were assigned using the same UNITE v9.0 classifier applied to the full-length ITS and ITS2 data. This dataset was used for sensitivity analyses of taxonomic assignment and community structure.

### Data Analysis

All statistical analyses and data visualizations were performed in R Studio (v2024.04.2). ITS2 sequencing depth sufficiency was evaluated using rarefaction curves and Good’s coverage (1 − singletons/total reads). To account for compositionality and differences in sequencing depth, ASV count data were centered log-ratio (CLR) transformed using the microbiome::transform function (35). Taxa were agglomerated to higher taxonomic ranks using the tax_glom function in phyloseq (36) as needed. Alpha-diversity metrics, including Shannon diversity and Chao1 richness, were calculated using vegan (37). Beta diversity was assessed using Bray-Curtis dissimilarity on relative abundance data and visualized with principal coordinate analysis (PCoA). Differences in community composition were tested using permutational multivariate analysis of variance (PERMANOVA) via adonis2 with 10,000 permutations, using Euclidean distances on CLR-transformed data and accounting for paired sample structure where applicable. Ordination concordance between datasets was evaluated using symmetric Procrustes analysis (PROTEST) and Mantel tests (Spearman) comparing Bray-Curtis distance matrices. Taxonomic assignment success between Illumina ITS2 and PacBio full-length ITS pipelines was quantified as the percentage of ASVs assigned at each taxonomic rank per paired sample. Genus-level abundance agreement between pipelines was assessed using Spearman correlations on CLR-transformed data. Differences in abundance of health-relevant genera between pipelines and sample types were tested using paired Wilcoxon tests. Differences in per-sample correlation coefficients across sample types were evaluated using analysis of variance (ANOVA) followed by Tukey’s HSD where appropriate. Assumptions of normality and homogeneity were assessed using Shapiro–Wilk and Levene’s tests.

## Results

### Comparison of ITS2 and Full-Length ITS Sequencing Pipeline Performance

PacBio full-length ITS sequencing yielded substantially higher post-QC read counts and ASV richness than ITS2 (mean = 739,601 ± 1,565,353 reads per sample; 6,419 ASVs), whereas Illumina ITS2 produced 10,395 ± 6,817 reads per sample and 1,814 ASVs. Illumina ITS2 libraries showed sufficient sequencing depth for downstream analyses, as indicated by rarefaction curves that approached saturation at ∼ 2k reads for nearly all samples (Fig. S2). Good’s coverage values were similarly high (median = 1.00; IQR = 1.00-1.00; mean = 0.97), confirming that most ITS2 samples were well covered. Three samples had low coverage (< 0.90) and were removed from paired ITS2-full-length ITS comparisons, taxonomic assignment analyses, and diversity tests; all descriptive summaries retained the full dataset. Sequence length profiles are shown in Fig. S3. Full-length ITS reads spanned ∼400-800 bp with a peak around 550 bp; ITS2 reads from Illumina pipeline or trimmed from full-length ITS sequences were similar (∼100-300 bp, peak ∼ 150-200 bp), reflecting the targeted subregion and potential trimming effects. We further compared both per-sample percentage of ASVs assigned and total number of ASVs classified at each taxonomic rank of two pipelines (Fig. 1a). Both pipelines showed similar performance across the higher ranks, with nearly complete assignment at kingdom (100%), phylum, class, and order, and comparable results at the genus level (75%). Differences emerged at finer ranks. Full-length ITS assigned 5,742 ASVs (98% of paired-sample mean) at the family level and 2,473 ASVs (42%) at the species level, whereas ITS2 assigned 1,354 ASVs (87%) at the family level and 679 ASVs (32%) at the species level.

**Fig 1.**
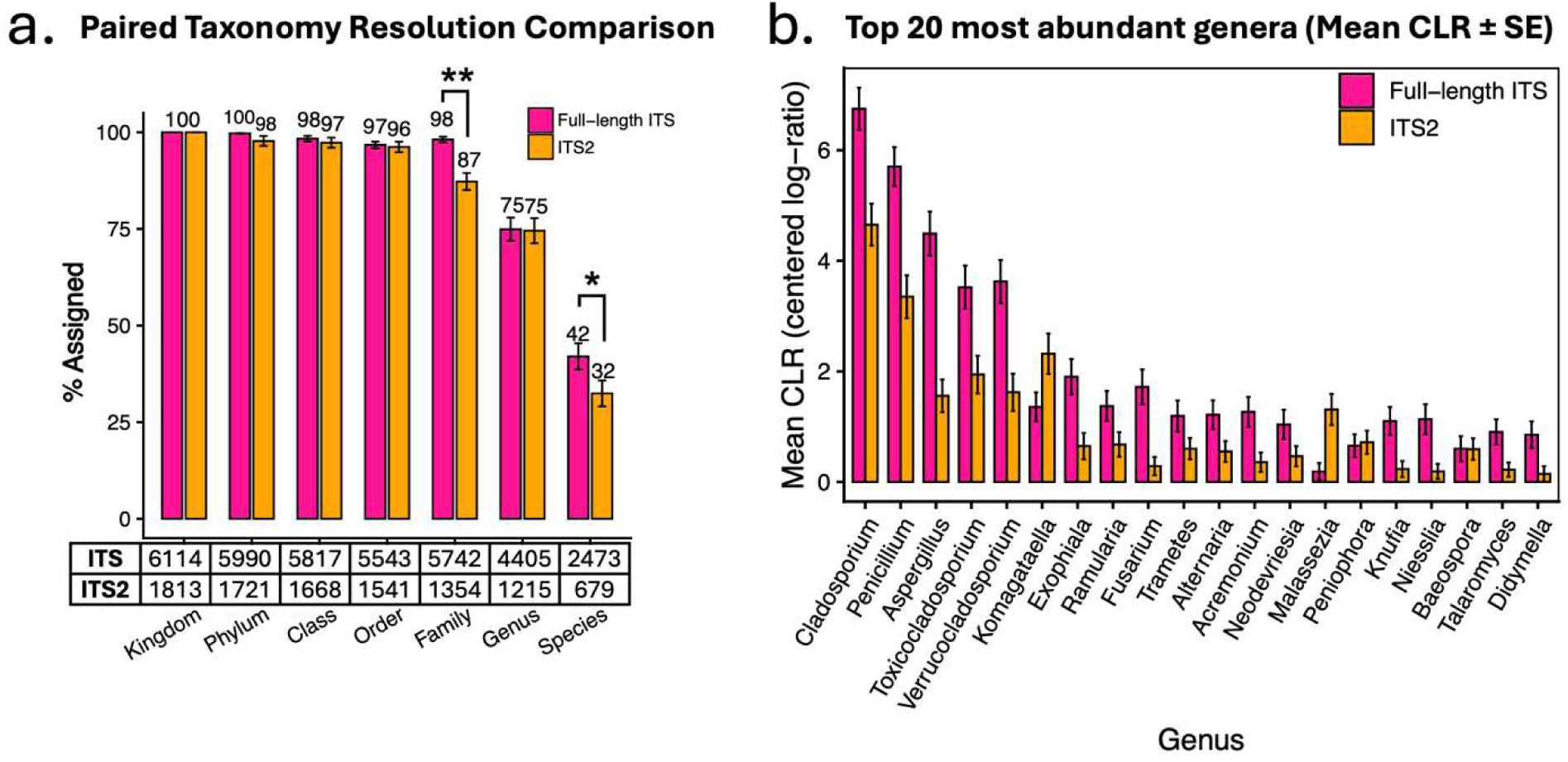
Comparative performance of Illumina ITS2 and PacBio full-length ITS sequencing platforms. (a) Taxonomic assignment across all ranks comparing PacBio full-length (deep pink) and Illumina ITS2 (orange) pipelines by paired sample comparison, numbers above the bars indicate percent of ASV classified ± standard error (SE) for each rank, numbers in the table under the plot showed the total number of ASVs classified at each rank from each pipeline. (b) Top 20 genera based on mean CLR abundance across both sequencing pipelines. Bars show mean centered log-ratio (CLR) abundance ± standard error (SE) for each genus, with paired values plotted for full-length ITS and ITS2.

At the phylum level (Fig. S4), both pipelines were dominated by Ascomycota and Basidiomycota. Chytridiomycota was detected only in the ITS2 dataset, reflecting our inclusion of chytrid-specific primer variants in the ITS2 pipeline improved amplification of these lineages. The top 20 most abundant genera based on mean CLR abundance showed broadly similar patterns between the two sequencing pipelines, particularly among the highest-abundance taxa (Fig. 1b). Both methods recovered *Cladosporium*, *Penicillium*, *Aspergillu*s, and *Toxicocladosporium* as dominant genera. Several taxa, including *Komagataella* and *Malassezia*, displayed higher CLR abundance in the ITS2 data, whereas *Didymella*, *Fusarium*, *Talaromyces*, and *Trametes* were more abundant in the full-length ITS dataset. Divergence increased among lower-abundance genera, where full-length ITS resolved several taxa that appeared reduced or absent in ITS2 profiles (Fig. S5). These differences likely reflect primer-specific amplification patterns.

### Community-Level Diversity and Compositional Differences Between Sequencing Pipelines

Paired comparisons of alpha diversity showed that the two pipelines produced similar overall diversity profiles across sample types (Fig. 2a). Shannon diversity did not differ between ITS2 and full-length ITS in any sample type (paired Wilcoxon, all q ≥ 0.05), and values ranged from 0 to 6.5. In contrast, richness metrics showed consistent increases for the full-length ITS dataset. Chao1 richness was significantly higher for full-length ITS in air, swab, and negative control samples (paired Wilcoxon q < 0.05). Paired scatter plots further showed weak sample-level correlation in Shannon diversity and poor concordance in richness estimates between pipelines (Fig. S6), despite similar group-level Shannon distributions and consistently higher richness recovered by full-length ITS.

**Fig 2.**
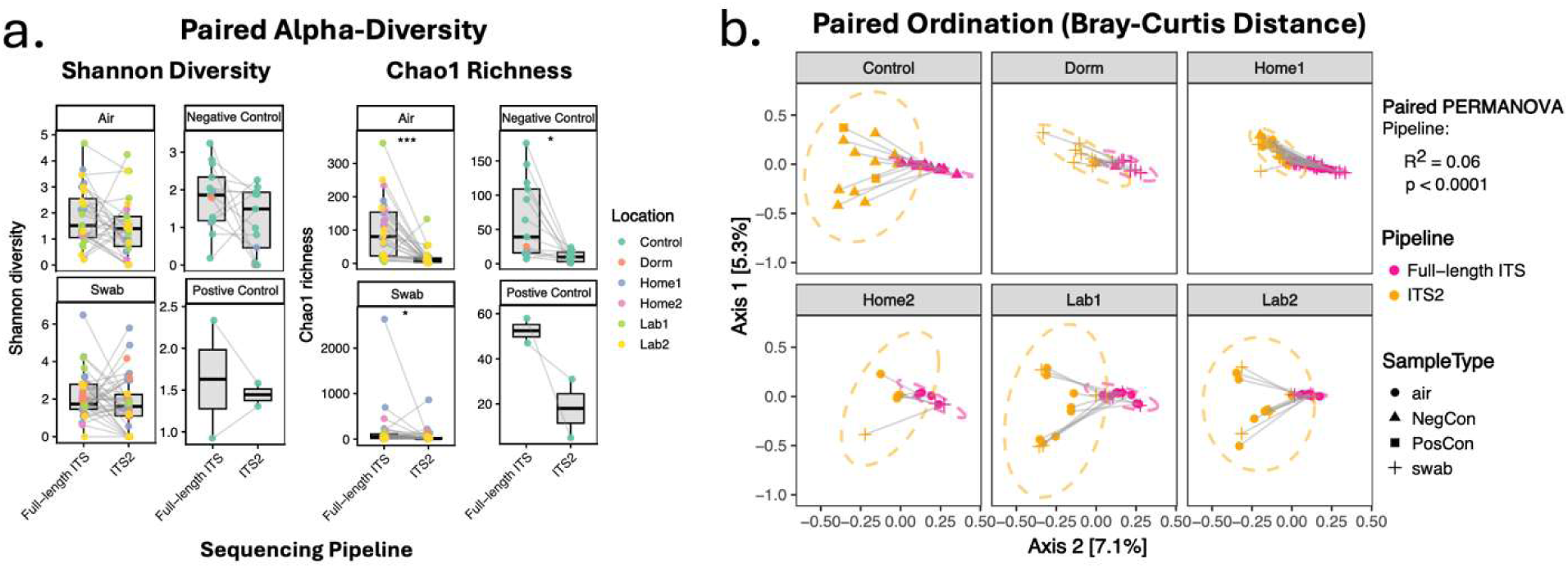
Community-Level Diversity and Compositional Differences Between Sequencing Pipelines. (a) Paired alpha-diversity comparisons for ITS2 and full-length ITS across sample types. Side-by-side boxplots show Shannon diversity (left) and Chao1 richness (right) for paired samples, with connecting lines linking the same sample across pipelines. Points are colored by sampling location. Statistical significance labeled for paired Wilcoxon test comparing alpha-diversity index values between pipelines. (b). Paired ordination comparing ITS2 and full-length ITS profiles using Bray-Curtis dissimilarity. Principal coordinates analysis (PCoA) shows each sample pair connected by a line, with ITS2 and full-length ITS placed in proximity but occupying distinct positions. Ellipses represent 95% confidence intervals for location groups. Paired PERMANOVA indicated a significant pipeline effect (R² ≈ 0.06, p < 0.001), consistent with the systematic shifts observed between ITS2 and full-length ITS profiles. Asterisks indicate statistically significant differences between short vs long reads or sample types (*** for p-value between 0-0.001, ** for p-value between 0.001-0.01, * for p-value between 0.01 - 0.05).

Paired and unpaired beta-diversity analyses showed that the two sequencing pipelines captured broadly similar community-level patterns, although full-length ITS revealed stronger structure associated with environmental variables than ITS2 (Fig. 2b; Fig. S8). In paired PERMANOVA restricted to matched samples, sequencing workflow explained a small but significant fraction of Bray-Curtis dissimilarity (R² ≈ 0.06, p < 0.0001), and paired ordination plots showed a consistent shift between ITS2 and full-length ITS profiles derived from the same samples (Fig. 2b).

Unpaired principal coordinates analysis showed clearer separation by sample type and location for the full-length ITS dataset than for ITS2 (Fig. S8). PERMANOVA indicated stronger effects of location in the full-length ITS dataset (R² = 0.12, p < 10⁻⁴) than in the ITS2 dataset (R² = 0.10, p < 10⁻⁴), with sample type contributing a smaller but significant proportion of explained variance in both cases. In contrast, the PacBio-derived in-silico ITS2 dataset displayed ordination patterns in between full-length ITS and Illumina ITS2 (location: R² = 0.11, p < 10⁻⁴), and similar sample type separation (R² = 0.06, p = 0.0001) to full-length ITS. Measures of cross-pipeline agreement supported this interpretation (Fig. S8). Mantel correlations between Bray-Curtis distance matrices were weak and not significant for both ITS2 versus full-length ITS (r = 0.06, p = 0.098) and ITS2 versus trimmed ITS2 (r = 0.03, p = 0.273). In contrast, symmetric Procrustes rotation of full ordination configurations indicated moderate and significant alignment in both comparisons (ITS2 vs full-length ITS: r = 0.67, p = 0.001; ITS2 vs trimmed ITS2: r = 0.63, p = 0.001). Restricting the analysis to the first 15 PCoA axes reduced alignment strength but retained statistical significance (r ≈ 0.40, p = 0.001), indicating that shared community structure is distributed across multiple ordination dimensions rather than confined to the dominant axes. These results implicate amplicon length as the primary driver of differences in community-level inference.

### Genus-Level Agreement and Taxon-Specific Differences Between Sequencing Pipelines

Agreement between sequencing pipelines was evaluated at the genus level (Fig. 3). Spearman correlations of CLR-transformed abundances for overlapping genera ranged from approximately 0 to 0.7 (mean ± SD: 0.27 ± 0.15) across sample types and locations (Fig. 3a). Correlation strength differed significantly by sample type. Abundance was significantly higher in air samples compared with positive control samples (ANOVA p = 0.001), indicating that the degree of cross-pipeline similarity depends on sample characteristics rather than solely on sequencing pipelines.

**Fig 3.**
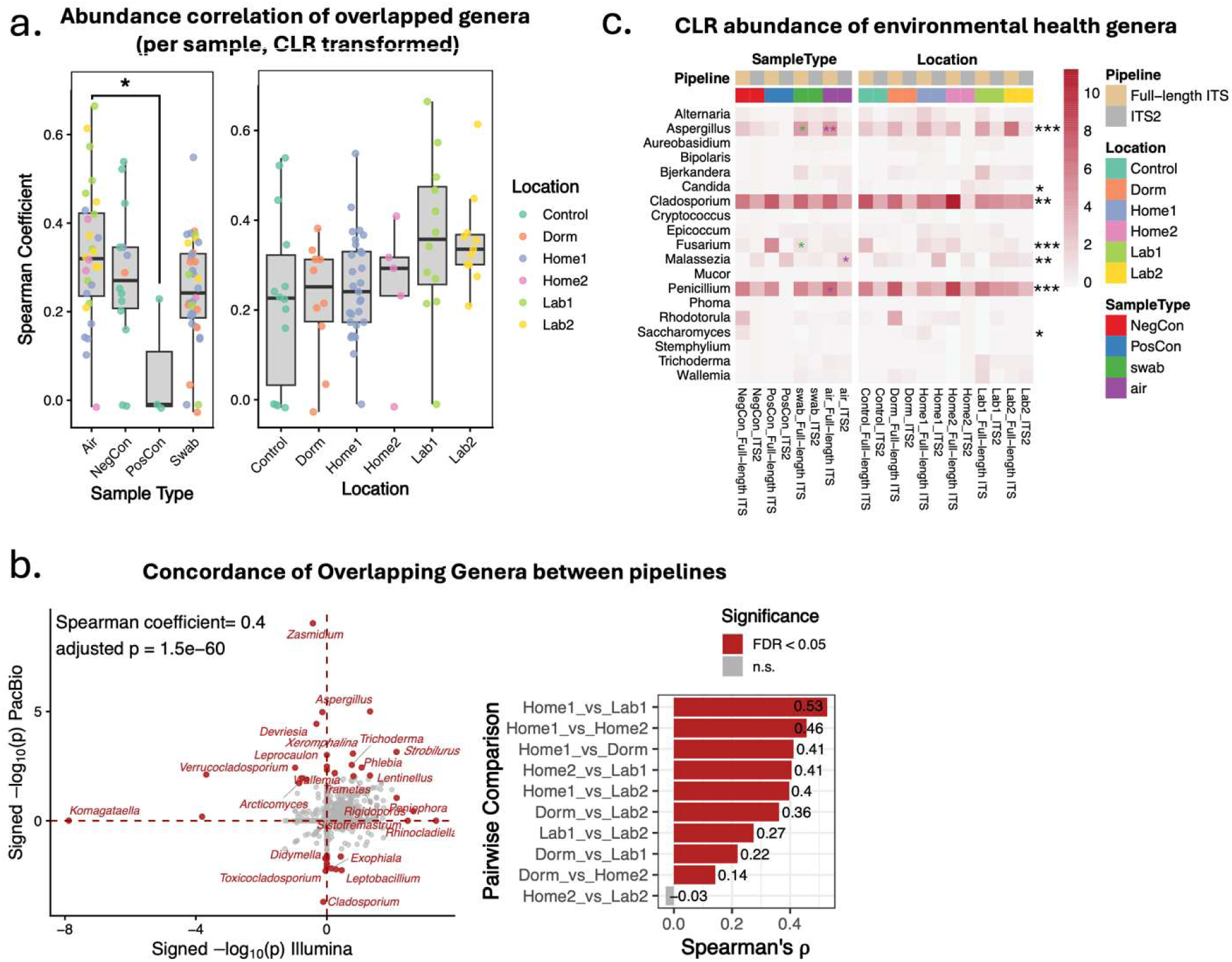
Genus-level agreement and taxon-specific differences between sequencing pipelines. (a) Abundance correlation of overlapping genera. Spearman correlations (per sample, CLR-transformed abundances) between ITS2 and full-length ITS for overlapping genera, grouped by sample type (left) and location (right). Correlation strength varied significantly across sample types and locations, with laboratory samples showing the highest cross-pipeline agreement. Each point represents one sample, colored by sampling location. (b) Concordance of taxon-level statistical inference across pipelines. Top: Scatter plot of signed −log₁₀(p-values) from differential abundance analyses for shared genera across all pairwise environmental contrasts, comparing ITS2 (x-axis) and full-length ITS (y-axis). Red points indicate genera with FDR < 0.05 in either pipeline. Dashed red lines mark direction reversals. Annotated genera highlight lineages with strong or pipeline-specific signals. The overall Spearman correlation (ρ = 0.40, adjusted p = 1.5×10⁻⁶⁰) indicates moderate global concordance. Right: Spearman correlations for each pairwise location comparison. Bars show correlation strength, with significant comparisons (FDR < 0.05) highlighted. c. Abundance patterns of health-relevant genera. Heatmap of mean CLR abundance for selected health-relevant fungal genera across sample types (air, swab, controls) and indoor locations (Home1, Home2, Dorm, Lab). Asterisks indicate significant differences between pipelines or sample types (*** p < 0.001, ** p < 0.01, * p < 0.05). Several genera displayed strong pipeline-dependent or location-specific patterns, illustrating how sequencing strategy and environmental context shape the detected indoor mycobiome.

To evaluate concordance in statistical inference across pipelines, we compared signed-log_10_ (p-values) from differential abundance tests for all shared genera and environmental contrasts (Fig. 3b, left). The plot indicated a moderate but highly significant correlation between pipelines (Spearman’s ρ = 0.40, *p* = 1.5 × 10^-60^). Most genera clustered near the origin, reflecting weak or nonsignificant differences between pipelines, whereas a subset of genera exhibited strong signals in one or both datasets. Notably, a few genera (e.g. *Komagataella*, *Zasmidium*, *Aspergillus*) deviated far from the 1:1 line, indicating lineage-specific sensitivity due to primer targeting differences. To summarize how concordance varies across environmental contrasts, we computed Spearman correlation coefficients for each pairwise location comparison (Fig. 3b, right). Correlation values ranged from weak or negative to moderately strong (ρ ≈ –0.2 to 0.53), with most comparisons showing statistically significant agreement after false-discovery-rate correction. Together, these analyses demonstrate that while the two pipelines produce broadly similar genus-level inferences, pipeline-specific differences for genera or environmental contrasts may influence downstream ecological interpretation.

Health-relevant fungal genera exhibited distinct abundance patterns across sequencing pipelines, sampling types, and locations (Fig. 3c and Table S3). Ubiquitous indoor genera such as *Cladosporium*, *Penicillium*, and *Aspergillus* were consistently detected with high CLR abundances that were significantly different between pipelines (Wilcoxon adjusted p < 0.01). Several other genera, including *Malassezia*, *Fusarium*, *Candida*, and *Saccharomyces*, also showed significant differences in CLR abundance between sequencing pipelines (adjusted p < 0.05). Several genera exhibited sample-type-dependent shifts between pipelines (Fig. 3c). In swab samples, the full-length ITS pipeline detected higher CLR abundances of *Aspergillus* and *Fusarium* (adjusted p < 0.05). In air samples, full-length ITS also yielded higher CLR abundances of *Aspergillus* and *Penicillium*, whereas ITS2 showed higher CLR abundance of *Malassezia* (adjusted p < 0.05). These results indicate that pipeline influences the apparent abundance of selected health-relevant taxa in a manner that depends on sample type.

### Cross-Platform Taxonomic Mapping: Unmapped ASVs

To assess the degree of complementarity between sequencing pipelines, ITS2 ASVs were mapped against the full-length ITS ASVs using BLAST, treating the full-length ITS dataset as a reference. A total of 1,136 ITS2 ASVs (63%) mapped to at least one full-length ITS ASV at ≥99% identity, corresponding to 2,731 full-length ITS ASVs (45%) that received at least one ITS2 match (Fig. 4a). The remaining 677 ITS2 ASVs (37%) and 3,383 full-length ITS ASVs (55%) lacked detectable counterparts under these criteria. Analysis at the 100% identity further illustrated this pattern (Figure 4b). Most ITS2 ASVs (n = 491) showed a one-to-one relationship with full-length ITS sequences, while a subset (n = 384) mapped to multiple (2–10) full-length ITS variants. Conversely, the majority of full-length ITS ASVs had one single ITS2 ASV mapping (n = 2,046), with very few having two ITS2 sequence mapping (n =7). Together, these results demonstrate that the two pipelines recover overlapping but non-identical representations of fungal diversity. Differences in mapping patterns, including one-to-many relationships at 100% identity, are consistent with amplicon-length-driven resolution differences and primer targeting effects, rather than the absence of taxa in either dataset.

**Fig 4.**
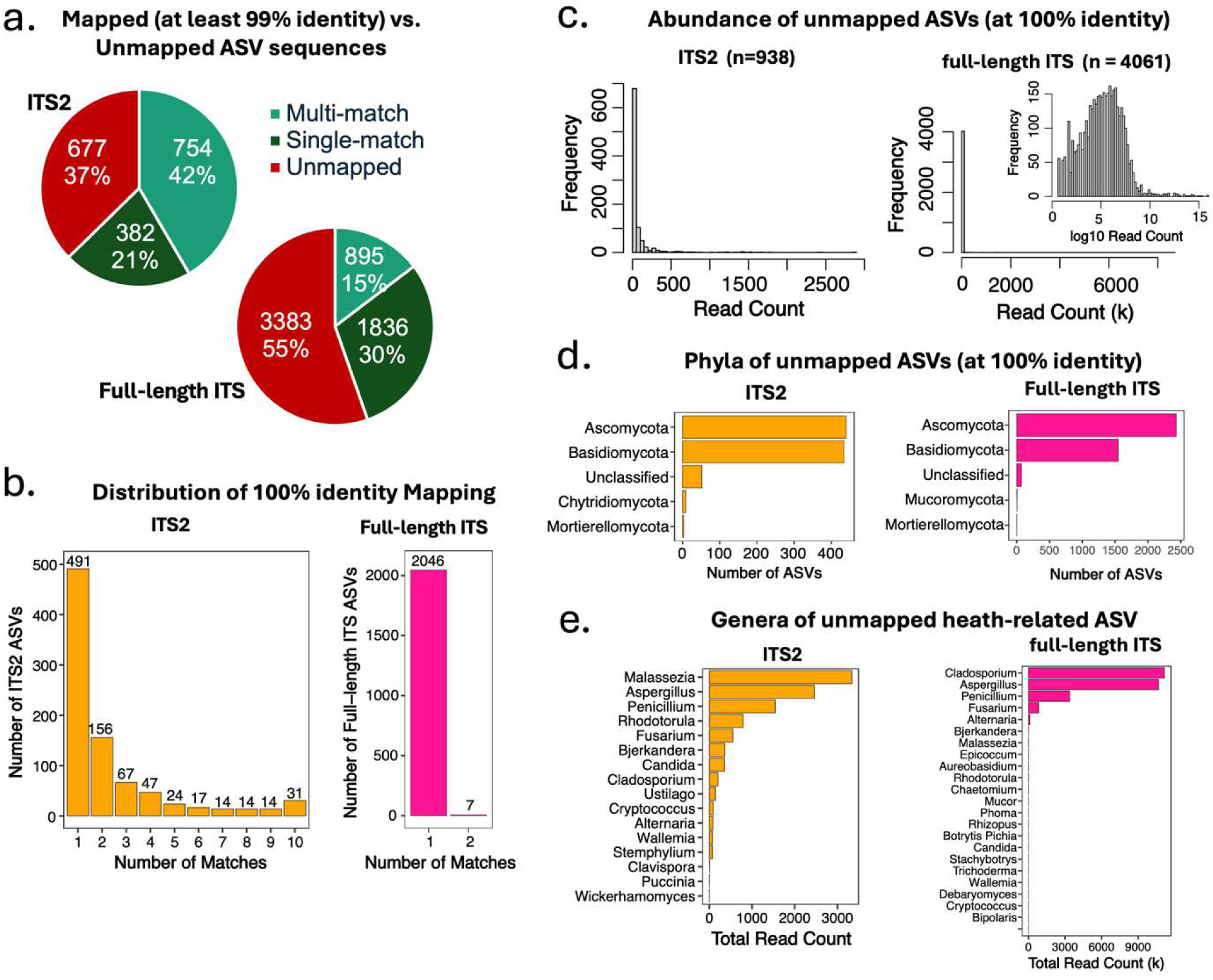
Cross-platform taxonomic mapping analysis of ITS2 pipeline against full-length ITS pipeline, using full-length ITS data as the reference dataset. (a) Pie chart showing mapped and unmapped ASV counts and percentages in ITS2 (left) and full-length ITS (right) pipelines. Unmapped indicates ITS2 ASV sequence was not mapped to any of full-length ITS sequence, single match indicates ITS2 sequence was mapped to one full-length ITS sequence, multi-match indicates ITS2 sequence mapped to 2-10 full length ITS sequence. (b) Distribution of 100% identity mapping. (c) Read abundance distribution of unmapped ITS2 (left) and full-length ITS (right) ASVs demonstrating predominance of low-abundance taxa. (d) Phyla distributions of unmapped ITS2 (left) and full-length ITS (right) ASVs. (e) Read abundance of unmapped ASVs assigned to health-related genera from the ITS2 (left) and full-length ITS (right) pipelines.

Unmapped ASVs from both pipelines were predominantly low in read abundance (Fig. 4 c-d). Most unmapped ITS2 ASVs had fewer than 500 total reads, with the majority below 100 reads (Fig. 4c). Although most unmapped full-length ITS ASVs also occurred at low abundance, a subset displayed moderate to high read counts when visualized on a log₁₀ scale (Fig. 4c, right). This increase likely reflects both the substantially higher sequencing depth of the full-length ITS dataset and the greater resolving power of long-read amplicons. As a result, biologically abundant lineages that are represented by a small number of ITS2 ASVs can appear as multiple unmapped full-length ITS ASVs under strict 100% identity mapping criteria. Unmapped ASVs from both pipelines were primarily affiliated with Ascomycota and Basidiomycota, with smaller contributions (<100 ASVs) from under-classified fungi, Chytridiomycota (ITS2), Mucoromycota (full-length ITS), and Mortierellomycota (both pipelines) (Fig. 4 d). A subset of unmapped ASVs belonged to genera with relevance to indoor environmental health (Fig. 4e). Unmapped ITS2 ASVs included low-abundance assignments to genera *Malassezia*, *Aspergillus*, *Candida*, *Penicillium*, and *Fusarium* (Fig. 4e, left). Unmapped full-length ITS ASVs also included genera *Cladosporium*, *Aspergillus*, *Penicillium*, and *Fusarium*, often with higher cumulative read counts than their ITS2 counterparts (Fig. 4e, right). These unmapped health-associated ASVs likely reflect the influences of primer targeting and amplicon length on pipeline-specific detection, which can vary even within clinically relevant lineages. Together, these results demonstrate that ITS2 and full-length ITS pipelines provide overlapping yet distinct representations of fungal community composition, with primer targeting, amplicon length, and sequencing depth contributing to pipeline-specific detection of low-abundance taxa.

### Cross-Platform Taxonomic Mapping: mapped ASVs

Comparison of phylogenetic trees constructed from mapped ASVs demonstrates the enhanced resolving power of long-read sequencing (Fig. 5a). Although both phylogenies were largely bifurcating, the ITS2-derived tree showed significantly shorter branch lengths than the full-length ITS tree (median 0.0266 vs. 0.0367; Wilcoxon test, p = 1.6 × 10⁻⁸, Fig. S9), indicating reduced phylogenetic signal in the ITS2 region. Longer branches in the full-length ITS tree reflect increased sequence variation across ITS1, 5.8S, and ITS2. Among the ITS2 ASVs mapped at 100% identity to full-length ITS sequences, taxonomic assignment discrepancies were observed at multiple ranks (Fig. 5b). A total of 17 ASVs had taxonomy assignment differed at the species level and 6 at the genus level. These discrepancies likely reflect the increased discriminatory power of full-length ITS sequences. Cross-platform comparison also revealed shifts in taxonomic resolution (Fig. 5c). Among the mapped ITS2 ASVs, the majority (599 ASVs) remained unchanged, suggesting broad agreement of taxa with robust representation across databases. Around 15% ASVs (n = 131) gained higher-resolution classification after mapping to full-length ITS reference, highlighting the added discriminatory information obtained from longer reads. Around 14% ASVs (n = 121) were reassigned to broader taxonomic ranks, likely due to the primer target differences and the limited coverage of full-length ITS reference we constructed. These patterns illustrate that cross-platform mapping exerts an uneven influence on taxonomic resolution, depending on both sequence informativeness and reference database completeness.

**Fig 5.**
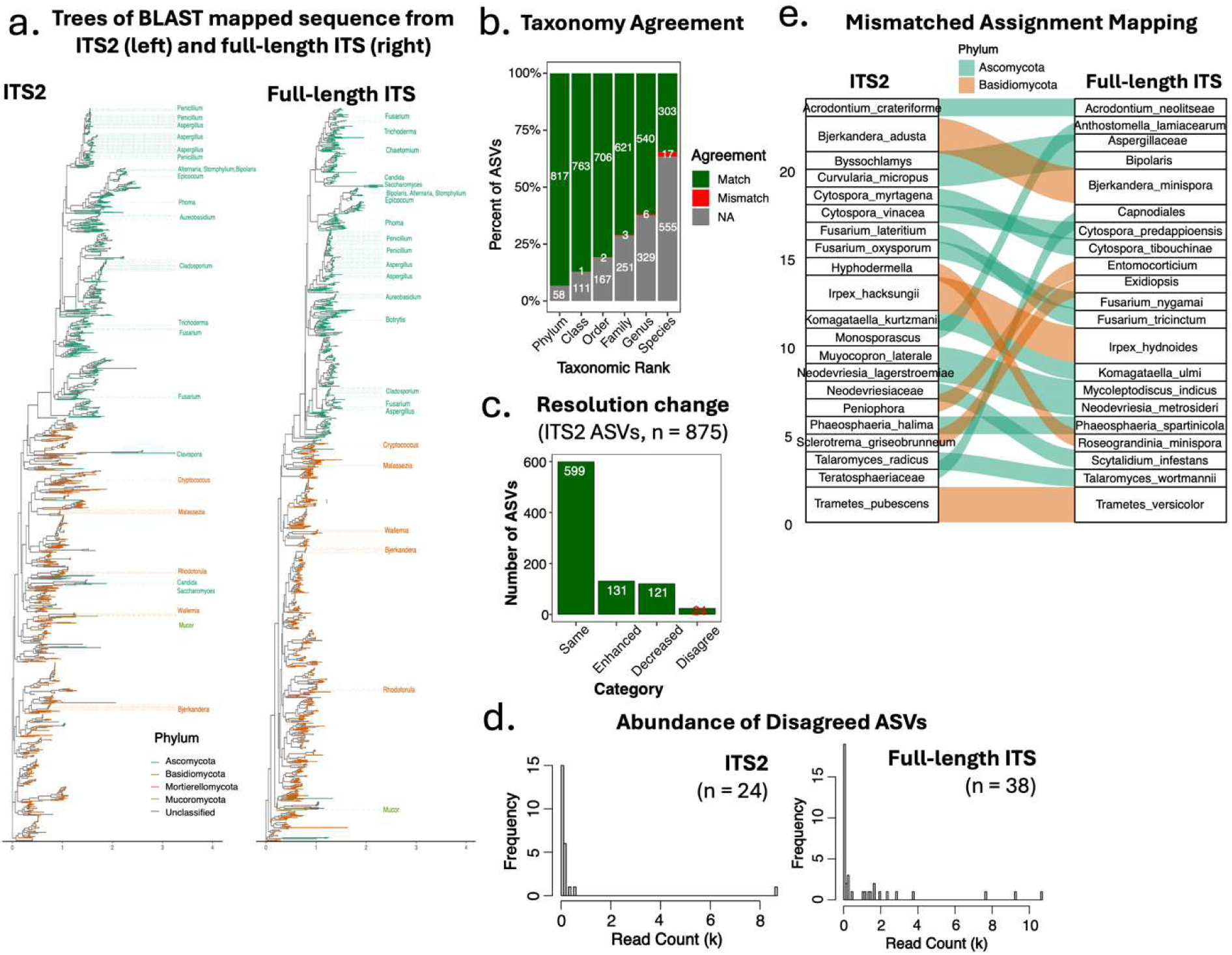
Cross-platform taxonomic mapping analysis of ITS2 pipeline against full-length ITS pipeline, using full-length ITS data as the reference dataset. (a) Rooted maximum likelihood trees constructed using BLAST mapped (> 97% identity) sequence from ITS2 (left) and full-length ITS (right) pipelines. The branches are colored by phylum, and health-related taxa are labeled at the genus level. (b) Taxonomic agreement analysis for 100% mapped ASVs, showing number of mismatched assignments at each taxonomic rank, NA(gray) indicates missing data in either or both pipelines. (c) Changes in ITS2 taxonomic resolution following mapping (100% identity), categorized as decreased, enhanced, or same resolution compared to original assignments. (d) Abundance distribution of 100% mapped but taxonomically mismatched ITS2 and full-length ITS ASVs. (e) Sankey diagram of taxonomic assignment mapping of taxonomically mismatched ASVs: ITS2 (left) and full-length ITS (right), with flow colors representing major fungal phyla.

The 24 ITS2 ASVs that had different assignment after mapping to the full-length ITS dataset were predominantly low in read abundance (< 1000 reads, Fig. 5d), except one ASV (ASV1382, assigned to *Komagataella kurtzmanii*) had over 8000 total reads. It was mapped at 100% identity to two full length ITS ASVs (ASV1411 and ASV6353, both assigned to *Komagataella ulmi*) at low abundance of 1061 total reads. This suggests that classification disagreement mainly affects rare taxa rather than dominant community members. A Sankey diagram visualizing taxonomic transitions between ITS2 and full-length ITS classifications further illustrate these dynamics (Fig. 5e). Most ASVs retained stable higher-level taxonomic placement, with a few ASVs gained different genus or species assignments, demonstrating how short-read and long-read sequences can yield divergent interpretations for certain taxa. These shifts emphasize the need for cautious cross-platform comparisons and highlight the importance of comprehensive, high-quality reference databases for achieving accurate taxonomic resolution in mycobiome studies.

## Discussion

This study provides a systematic evaluation of Illumina ITS2 and PacBio HiFi full-length ITS sequencing for indoor fungal community profiling, with emphasis on how sequencing strategy influences taxonomic resolution, community-level inference, and detection of health-relevant taxa. By integrating paired analyses, in-silico ITS2 trimming, and cross-platform sequence mapping, we demonstrate that apparent pipeline differences arise primarily from amplicon length and primer targets (19, 20, 38, 39).

### Amplicon length as the dominant driver of taxonomic resolution

Full-length ITS assigned over 40% of ASVs to species compared with approximately 32% for ITS2, reflecting the additional sequence variation captured across ITS1, 5.8S, and ITS2. This improvement is consistent with prior work showing that short-amplicon barcoding limits discrimination among closely related taxa, including clinically relevant species that differ by few substitutions within ITS2 alone or have identical ITS2 regions (18, 20, 21), and parallels gains reported when full-length 16S rRNA sequencing is applied to bacterial communities (40, 41).

Reference database completeness imposes an additional constraint on both pipelines: the UNITE database remains incomplete for many indoor-relevant fungal lineages, potentially underestimating the resolution advantage of full-length ITS and introducing differential classification biases (18, 33). Critically, in-silico trimming of PacBio reads to ITS2 caused convergence toward Illumina ITS2 in both assignment rates (Fig. S7) and community structure (Fig. S8), confirming that amplicon length rather than platform-specific error models or sequencing depth underlies the observed differences (19, 38).

### Community-level diversity and structure are broadly conserved across pipelines

Community-level diversity metrics were broadly conserved despite differences in resolution. Shannon diversity did not differ between pipelines in any sample type, indicating that dominant taxa and overall evenness are robust to sequencing strategy, a pattern consistent with platform comparisons in bacterial microbiome research (40, 41). In contrast, richness metrics (Chao1 and observed ASVs) were consistently higher for full-length ITS, reflecting increased resolution of intra-taxon sequence variation that accumulates as elevated richness in long-read datasets (19, 20). At the beta-diversity level, both pipelines recovered broadly similar community patterns, but full-length ITS revealed stronger structure associated with sampling location and sample type. PERMANOVA indicated that workflow explained only a small fraction of compositional variance (R² ≈ 0.06) relative to environmental factors, Procrustes analyses showed moderate ordination alignment, and weak Mantel correlations indicated that cross-pipeline agreement is distributed across many dimensions rather than concentrated in primary axes. The intermediate ordination position of in-silico trimmed PacBio-ITS2 data between full-length ITS and Illumina ITS2 further corroborates amplicon length as the primary driver (38), and confirms that ITS2 captures major community gradients in indoor environments (7, 16) while full-length ITS enhances discrimination along secondary axes of environmental heterogeneity.

### Genus-level agreement and context-dependent discrepancies

Genus-level abundance patterns were generally concordant between pipelines, particularly for dominant indoor genera such as *Cladosporium*, *Penicillium*, and *Aspergillus*, consistent with their prevalence across sequencing-based indoor surveys worldwide (5–7, 42). However, agreement varied by sample type and environment, with laboratory samples showing higher cross-platform correlations than residential or dormitory environments. This context dependence likely reflects interactions between community complexity and primer performance. In occupied buildings, higher α-diversity and greater representation of amplification-resistant lineages may amplify primer-targeting biases documented for ITS3/ITS4-type primers, particularly for Basidiomycota (27, 39). Differential abundance testing showed moderate global concordance (Spearman ρ = 0.40) alongside lineage-specific discrepancies, with certain genera (including *Komagataella*, *Zasmidium*, and *Aspergillus*) deviating markedly from the 1:1 line. For example, a high-abundance ITS2 ASV assigned to *K. kurtzmanii* mapped at 100% identity to two full-length ITS ASVs assigned to *K. ulmi*, illustrating that species-level inference can diverge between pipelines even for abundant taxa. Researchers should therefore be cautious when cross-comparing differential abundance results across sequencing strategies (18, 43). Although both datasets were classified using the same reference database and algorithm, concordance remains limited by incomplete fungal ITS reference resources and the insufficient discriminatory power of ITS1 or ITS2 alone for many taxa (43). Consequently, long-read ITS can resolve sequence variation that remains ambiguous in shorter ITS2 queries, producing marker-dependent differences that reflect reference limitations rather than pipeline error.

### Health-relevant taxa are differentially affected by sample type and marker choice

Differences in health-relevant genera between pipelines were driven primarily by sample type rather than location. Full-length ITS detected higher CLR abundances of *Aspergillus* and *Fusarium* in swab samples, and of *Aspergillus* and *Penicillium* in air samples, while ITS2 showed higher apparent abundance of *Malassezia* in air samples. The enrichment of *Malassezia* in ITS2 data reflects its unusually short ITS regions, which are preferentially amplified during PCR, combined with strong UNITE database representation (39, 44). Conversely, the greater sensitivity of full-length ITS for *Aspergillus* and *Penicillium* likely reflects improved resolution of species complexes collapsed to single ITS2 ASVs (10, 11); these genera encompass clinically distinct species with direct implications for indoor exposure and risk assessment (6, 9).

### Cross-platform mapping reveals resolution, not detection, differences

Cross-platform mapping analyses reinforce these interpretations: while most ITS2 ASVs mapped to full-length ITS at ≥99% identity, full-length ITS frequently partitioned abundant taxa into multiple closely related ASVs that collapsed into single ITS2 sequences, consistent with ITS2 underestimating within-taxon sequence diversity at the ASV level (18, 19, 38). Unmapped ASVs from both pipelines were predominantly low-abundance, though a subset of unmapped full-length ITS ASVs had moderate read counts, reflecting higher PacBio sequencing depth and greater resolving power rather than detection failure. Notably, health-relevant genera including *Aspergillus*, *Penicillium*, and *Fusarium* appeared among unmapped ASVs in both datasets (9–11), highlighting the value of inspecting unmapped fractions in pipeline comparisons. Phylogenetic trees from mapped ASVs further illustrated resolution differences: full-length ITS trees exhibited longer branches and more bifurcating nodes than ITS2 trees at the same evolutionary scale (19, 21), and taxonomic discrepancies among 100%-identity-mapped ASVs at species and genus levels reinforce calls for comprehensive reference databases and transparent reporting of taxonomic resolution (18, 33, 43).

### Implications for indoor mycobiome studies

Taken together, these results support a complementary, use-case-driven approach to indoor mycobiome sequencing. ITS2 remains efficient and cost-effective for large-scale surveys and community-level comparisons where patterns among dominant taxa are of primary interest, and is well-suited to epidemiological studies linking indoor fungi to health outcomes (3, 5, 9). Full-length ITS is particularly valuable when species-level resolution informs exposure or clinical risk assessment, especially for genera such as *Aspergillus* and *Fusarium* (6, 10). Recent improvements in PacBio HiFi throughput via the Revio system and Kinnex library preparation have reduced per-sample costs and expanded multiplexing feasibility (22, 28). With these improvements, full-length ITS will become increasingly tractable for large built-environmental studies. Meanwhile, a hybrid strategy combining ITS2 for community-wide profiling with targeted full-length ITS validation of taxa of health concern provides a practical middle ground. Improved species-level identification also enables downstream interpretation relevant to human health. In particular, genomic and phenotypic markers associated with mycotoxin production, β-glucan content, immune activation, and other pathogenic traits can inform the development of targeted assays and more informative exposure and health-risk assessments (45–47). More broadly, we found that community complexity in occupied settings amplifies discrepancies between pipelines, highlighting the importance of transparent reporting of sequencing strategy, reference database version, and bioinformatic pipeline choices for meaningful cross-study comparisons (18, 27, 43), particularly given the documented geographic and environmental structuring of indoor fungal communities (7, 15, 16).

### Limitations

This study has several limitations. The number of sites was limited, which may restrict generalizability of environment-specific conclusions (7, 16). Taxonomic assignments depend on the completeness of database, which remains incomplete for many lineages and may differentially affect short- and long-read annotations (18, 33). In-silico ITS2 trimming does not fully recapitulate PCR amplification with ITS2-specific primers, including primer binding efficiency, amplicon length bias, and chimera formation (19, 38). In addition, this analysis relied on DNA-based profiling and did not assess fungal viability, activity, or mycotoxin production; future integration of RNA-based or culture-based approaches would provide a more complete picture of active indoor fungal communities (6, 9, 43, 48). Finally, the higher sequencing depth of the PacBio dataset may have contributed to differences in low-abundance taxon detection. Future studies should target more comparable sequencing depths across platforms to better disentangle depth effects from those of amplicon length and primer targeting. As whole-genome sequencing of environmental and built-environment fungal isolates expands, reference databases are expected to better support long-read ITS classification, further improving species-level resolution beyond what is currently achievable.

In conclusion, amplicon length is the primary determinant of differences between Illumina ITS2 and PacBio full-length ITS pipelines for indoor mycobiome analysis. Both pipelines recover broadly similar community-level patterns for dominant taxa, but full-length ITS provides enhanced taxonomic resolution, improved richness estimates, and stronger ecological discrimination at the family and species levels. Platform-specific differences in the apparent abundance of health-relevant genera including *Aspergillus*, *Penicillium*, *Fusarium*, and *Malassezia* highlight how sequencing strategy and sampling matrix jointly shape the interpretation of indoor fungal exposure. Method selection should be guided by study scale, required taxonomic resolution, and research or exposure-assessment objectives in built-environment mycobiome studies.

## Data Availability

Sequences have been deposited at NCBI SRA (National Center for Biotechnology Information Sequence Read Archive, https://www.ncbi.nlm.nih.gov/sra) under BioProject ID PRJNA1294855.

## Acknowledgement

This work was supported by the National Science Foundation Engineering Research Center Precision Microbiome Engineering (NSF PreMiEr ERC) through Award No. 2133504.

Sequencing services were provided by the Duke Microbiome Sequencing Core, and we thank the core staff for their technical support and guidance.

## Author Contributions

R.N. and M.D. conceptualized the study. M.D., R.N., A.A.F., J.N., R.J.V., J.W.A., Y.J.K. generated the methodology. D.B., M.L., X.L., T.C., S.P.C., H.H., and M.D. performed the investigation. M.D. conducted formal analysis. M.D., M.L., and S.P.C. wrote the original draft of the manuscript. M.D. performed the visualization. R.N., L.D., and M.D. acquired funding. J.N., R.J.V., A.A.F., J.W.A., and R.N. provided resources. R.N. supervised the work. All co-authors reviewed and edited the manuscript.

## Supplementary Figures

**Fig. S1.**
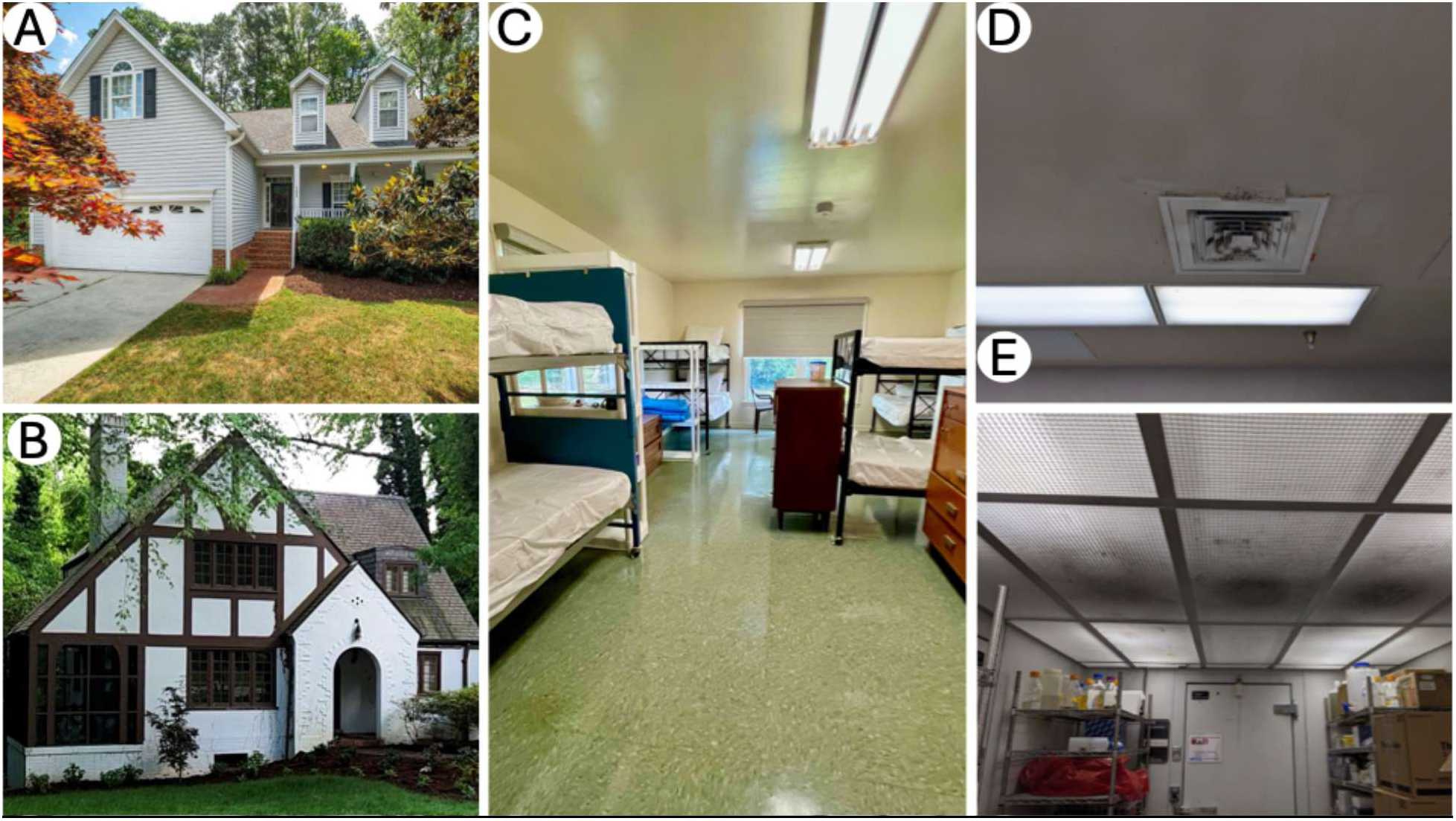
Images of sampling locations. (A) Home 1, an unoccupied residence in Carrboro, NC. (B) Home 2, a testbed residence for indoor environmental research at Duke University. (C) A student dormitory room at the UNC Institute of Marine Sciences (IMS). (D) Laboratory ceiling at the University of North Carolina at Chapel Hill (UNC), showing mold growth around an air vent. (E) Walk-in cold room at UNC with visible mold contamination on ceiling light diffuser panels.

**Fig S2.**
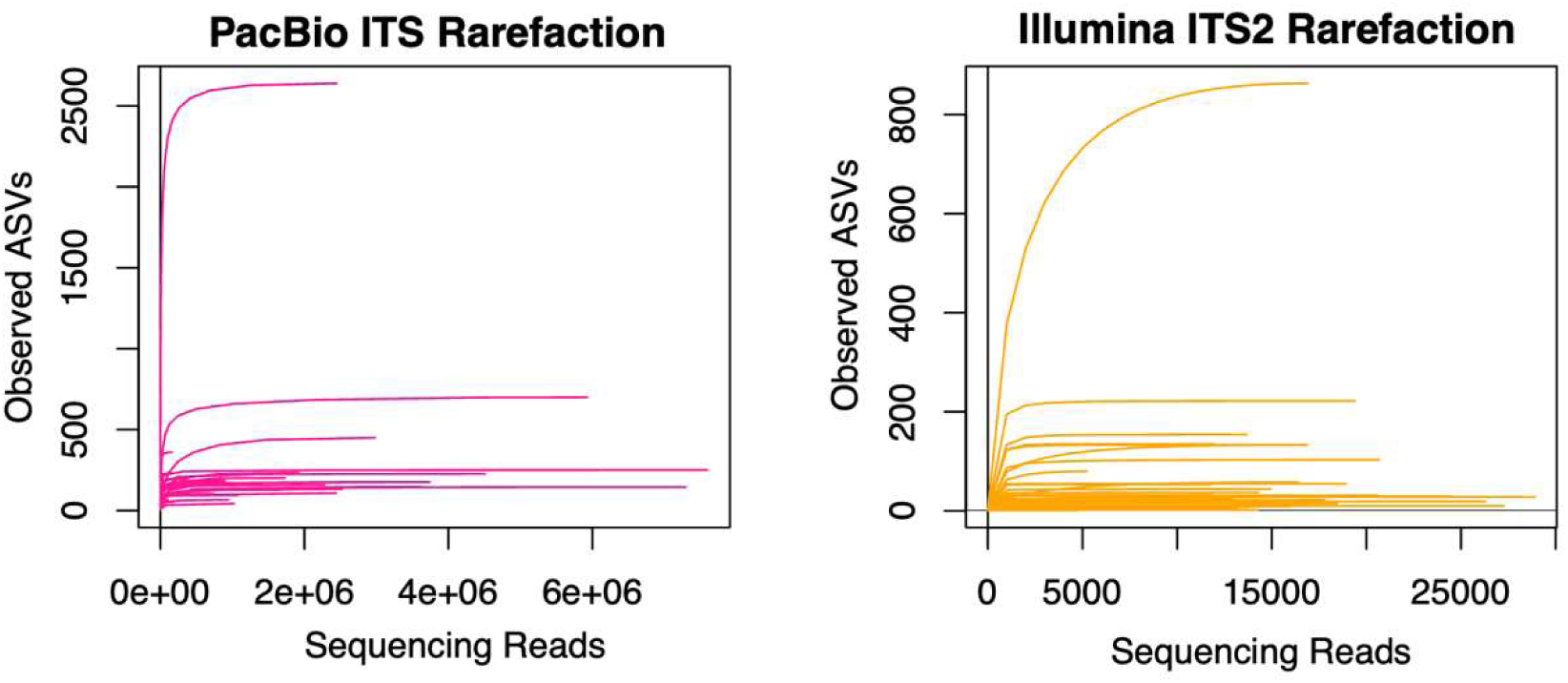
Rarefaction curves of full-length ITS (left) dataset and ITS2 (right) dataset.

**Fig S3.**
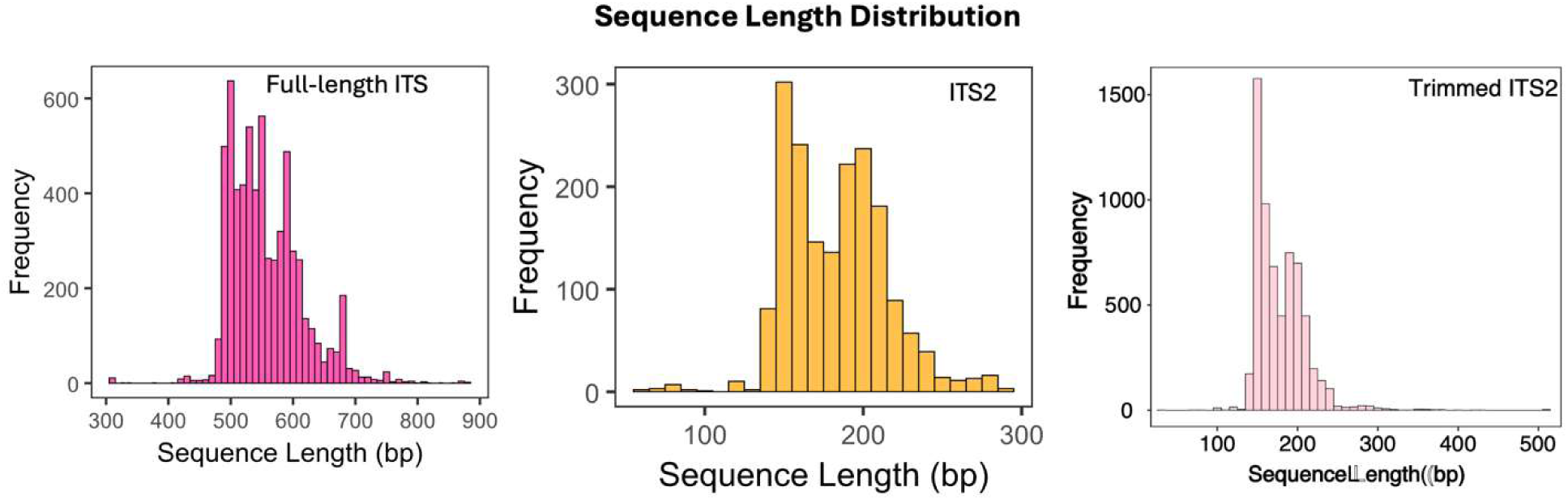
Sequence length distributions demonstrating the expected difference between full-length ITS sequences (upper panel, 400-800 bp), ITS2 fragments (middle panel, 100-300 bp), and in-silico trimmed ITS2 fragments (bottom panel, 100-300 bp).

**Fig. S4.**
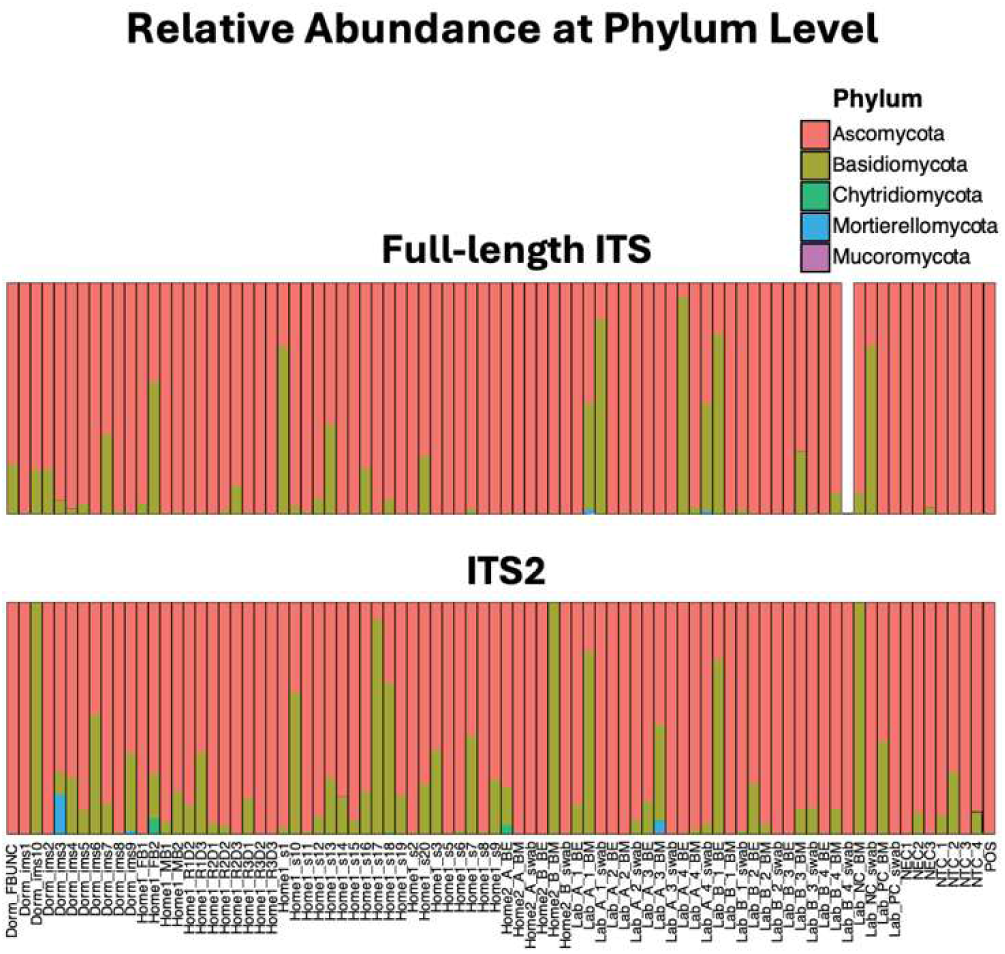
Relative abundance at phylum level derived from the full-length ITS pipeline (top panel) and ITS2 pipeline (bottom panel).

**Fig. S5.**
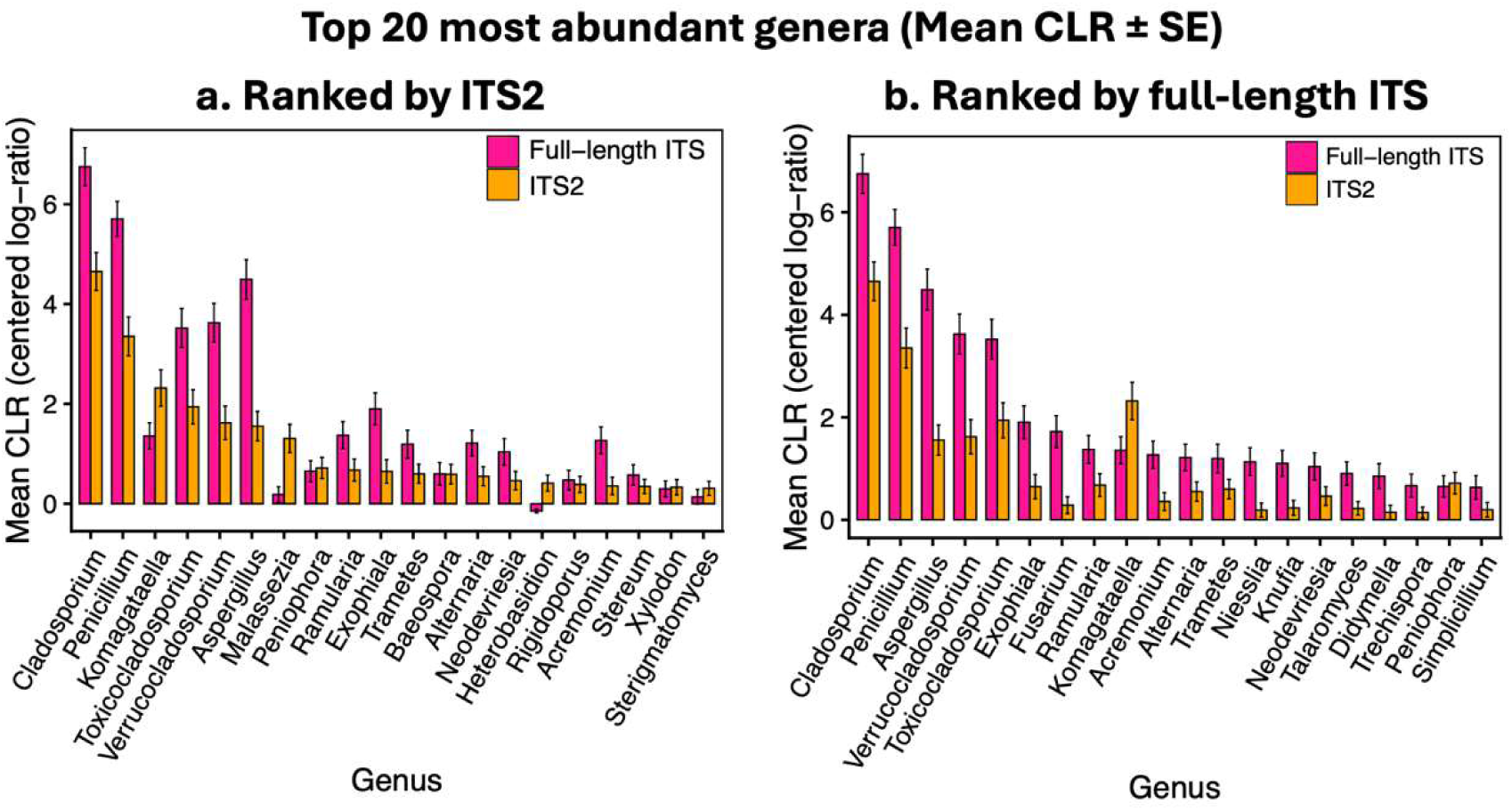
Top 20 genera ranked within each pipeline. (a) Top 20 by ITS2 mean CLR abundance; (b) Top 20 by Full-length ITS mean CLR abundance. Bars show mean CLR ± SE across samples. These panels illustrate method-specific rank shifts that are partially obscured when genera are ranked by combined abundance.

**Fig. S6.**
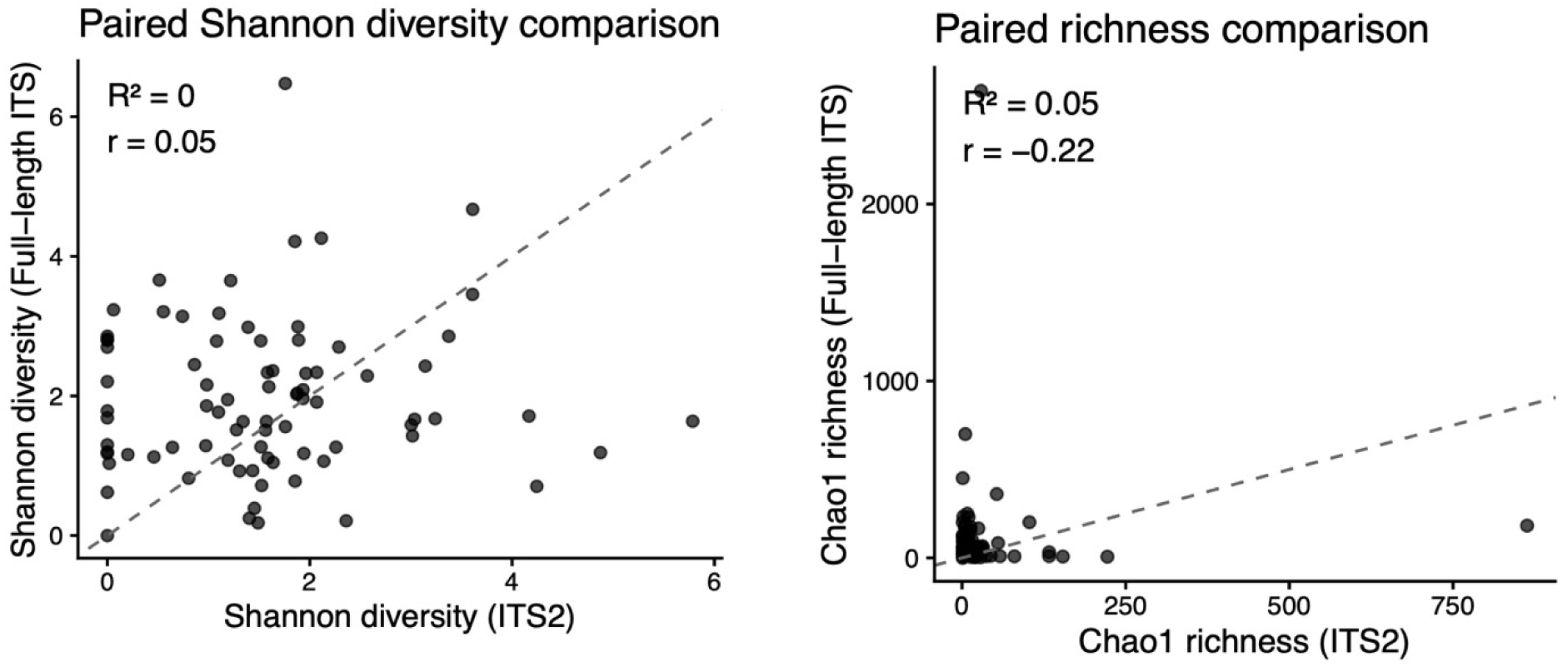
Paired comparisons of alpha-diversity metrics between sequencing pipelines. Paired scatter plots compare per-sample **Shannon diversity** (left) and **Chao1 richness** (right) estimated from Illumina ITS2 and PacBio full-length ITS datasets. Each point represents a matched sample processed by both pipelines, with dashed lines indicating the 1:1 relationship. Spearman correlation was used for both indexes. Shannon diversity showed minimal sample-level correlation despite similar group-level distributions, whereas richness estimates exhibited poor concordance, reflecting systematic differences in rare-taxon resolution across markers. These patterns indicate that alpha-diversity indices, particularly richness, are not directly interchangeable between ITS2 and full-length ITS pipelines.

**Fig. S7.**
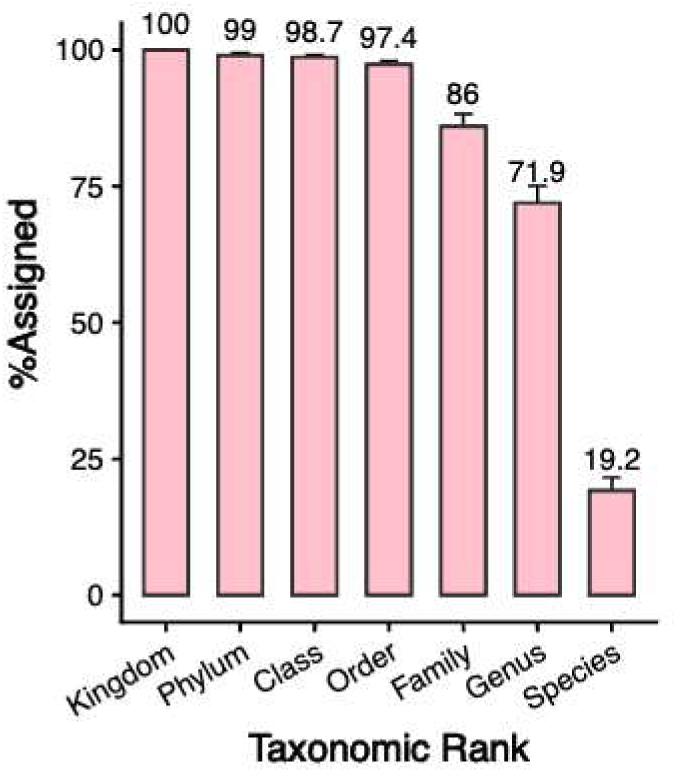
Taxonomic assignment across all ranks comparing of in-silico trimmed ITS2 sequences by paired samples. Numbers above the bars indicate percent of ASV classified ± standard error (SE) for each rank.

**Fig. S8.**
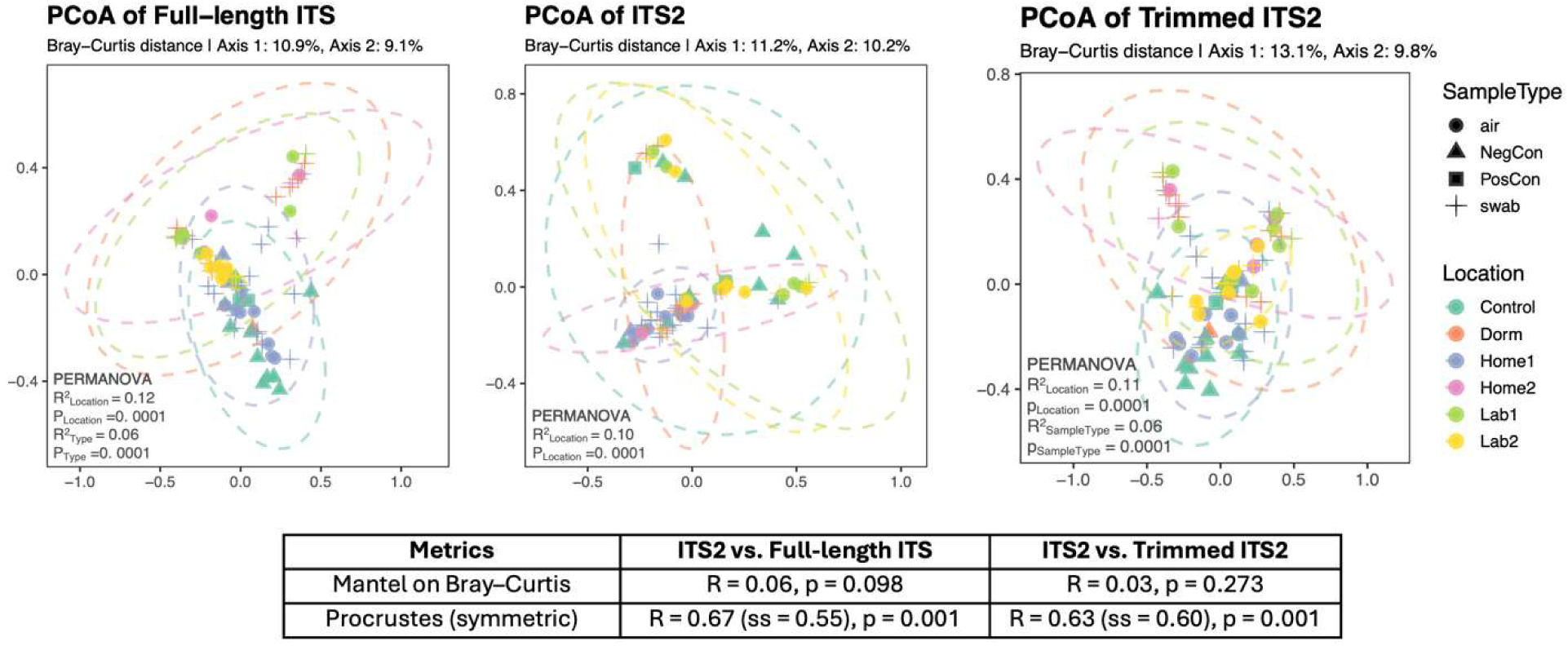
Comparison of full-length ITS, ITS2, and trimmed ITS2 ordinations based on Bray–Curtis dissimilarity. Principal coordinates analysis (PCoA) plots show fungal community composition derived from **full-length ITS** (left), **Illumina ITS2** (middle), and **PacBio-derived in-silico ITS2** (right). Ellipses represent 95% confidence intervals for location groups. Full-length ITS showed the strongest location-level separation (PERMANOVA R² = 0.12, p = 10⁻⁴), whereas both ITS2 and trimmed ITS2 displayed weaker but significant location structure (R² = 0.10, p = 10⁻⁴; and R² = 0.11, p = 10⁻⁴, respectively). Mantel tests comparing Bray–Curtis distance matrices indicated weak and non-significant correlations between ITS2 and full-length ITS (r = 0.06, p = 0.098) as well as between ITS2 and trimmed ITS2 (r = 0.03, p = 0.273). In contrast, symmetric Procrustes analysis showed moderate and significant alignment of ordination configurations for both comparisons (ITS2 vs full-length ITS: r = 0.67, sum of squares = 0.55, p = 0.001; ITS2 vs trimmed ITS2: r = 0.63, sum of squares = 0.60, p = 0.001). A robustness analysis restricted to the first 15 PCoA axes yielded lower but still significant alignment (r ≈ 0.40, p = 0.001), indicating that shared community structure is distributed across multiple ordination dimensions.

**Fig. S9.**
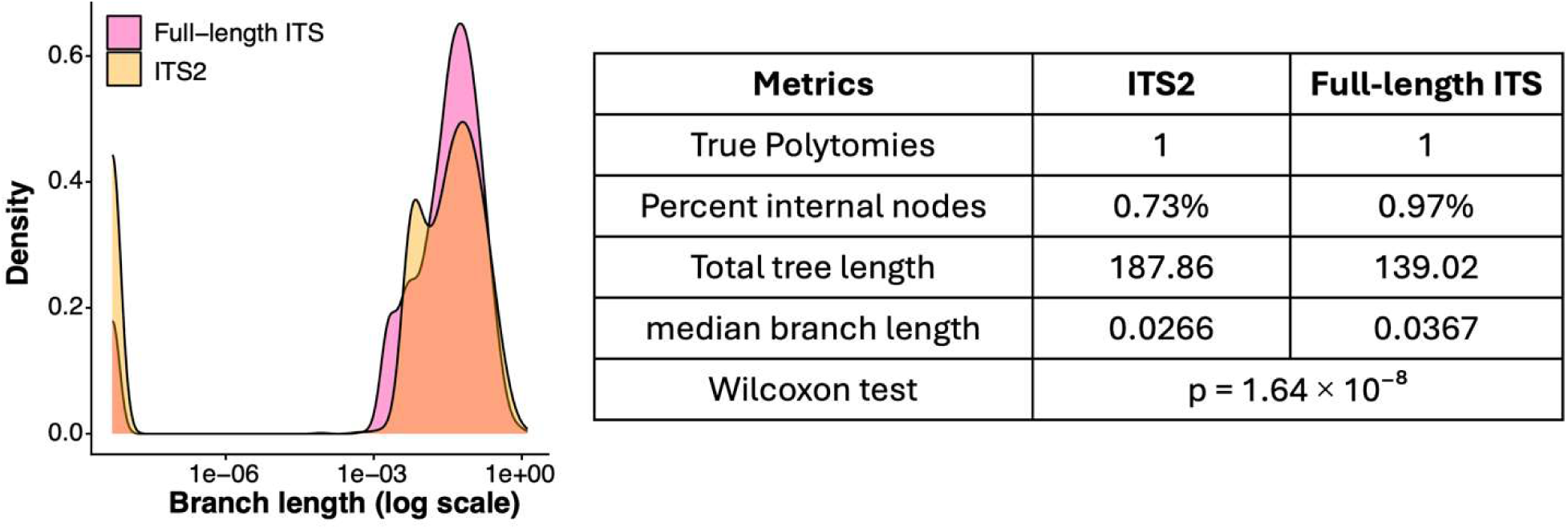
Branch length distributions for ITS2 and full-length ITS phylogenies. Kernel density plots show the distribution of branch lengths inferred from phylogenies constructed using ITS2 sequences (orange) and full-length ITS sequences (deep pink), plotted on a log scale. Both phylogenies were largely bifurcating, with a similarly low proportion of multifurcating internal nodes (polytomies; <0.1% of internal nodes in both trees). In contrast, branch lengths were significantly shorter in the ITS2-derived tree than in the full-length ITS tree (median branch length: 0.0266 vs. 0.0367; Wilcoxon rank-sum test, p = 1.64 × 10⁻⁸), indicating reduced phylogenetic signal in the ITS2 region. Longer branches in the full-length ITS phylogeny reflect greater sequence variation captured across the combined ITS1, 5.8S, and ITS2 regions. Although the ITS2 phylogeny exhibited greater total tree length, this reflected the accumulation of many short branches rather than increased phylogenetic resolution. In contrast, longer median branch lengths in the full-length ITS tree indicate greater information content per split.

## Supplementary Tables

**Table S1.** Samples summary.

| **Location** | **Sample (n)** | **Illumina ITS2**<br>**reads** | **PacBio full-length ITS**<br>**reads** |
| --- | --- | --- | --- |
| Home1<br>(n = 31) | Surface (19) | 195,294 | 15,429,812 |
|  | Air (8) | 80,283 | 4,203,379 |
|  | NegCon-Field Blank (2) | 43,224 | 3,613 |
|  | NegCon-Method Blank (2) | 16,903 | 3,640 |
| Home2<br>(n = 6) | Surface (2) | 26,610 | 4,727,465 |
|  | Air (4) | 45,776 | 8,336,048 |
| Dorm<br>(n = 11) | Surface (10) | 114,889 | 1,937,575 |
|  | NegCon-Field Blank (1) | 20,688 | 10,311 |
| Lab 1<br>(n = 12) | Surface (4) | 40,678 | 243,014 |
|  | Air (8) | 78,524 | 209,857 |
|  | NegCon-Method Blank (2) | 2,341 | 45,176 |
| Lab 2<br>(n = 12) | Surface (4) | 43,248 | 8,779 |
|  | Air (8) | 111,713 | 9,077,577 |
| Pipeline Control<br>(n = 12) | Positive Control (3) | 41,957 | 4,558,955 |
|  | NegCon -No Template (10) | 42,207 | 14,810,490 |

\*Reads reflect total raw reads in corresponding categories and were not adjusted for polling factors.
NegCon stands for Negative Control.

**Table S2.**
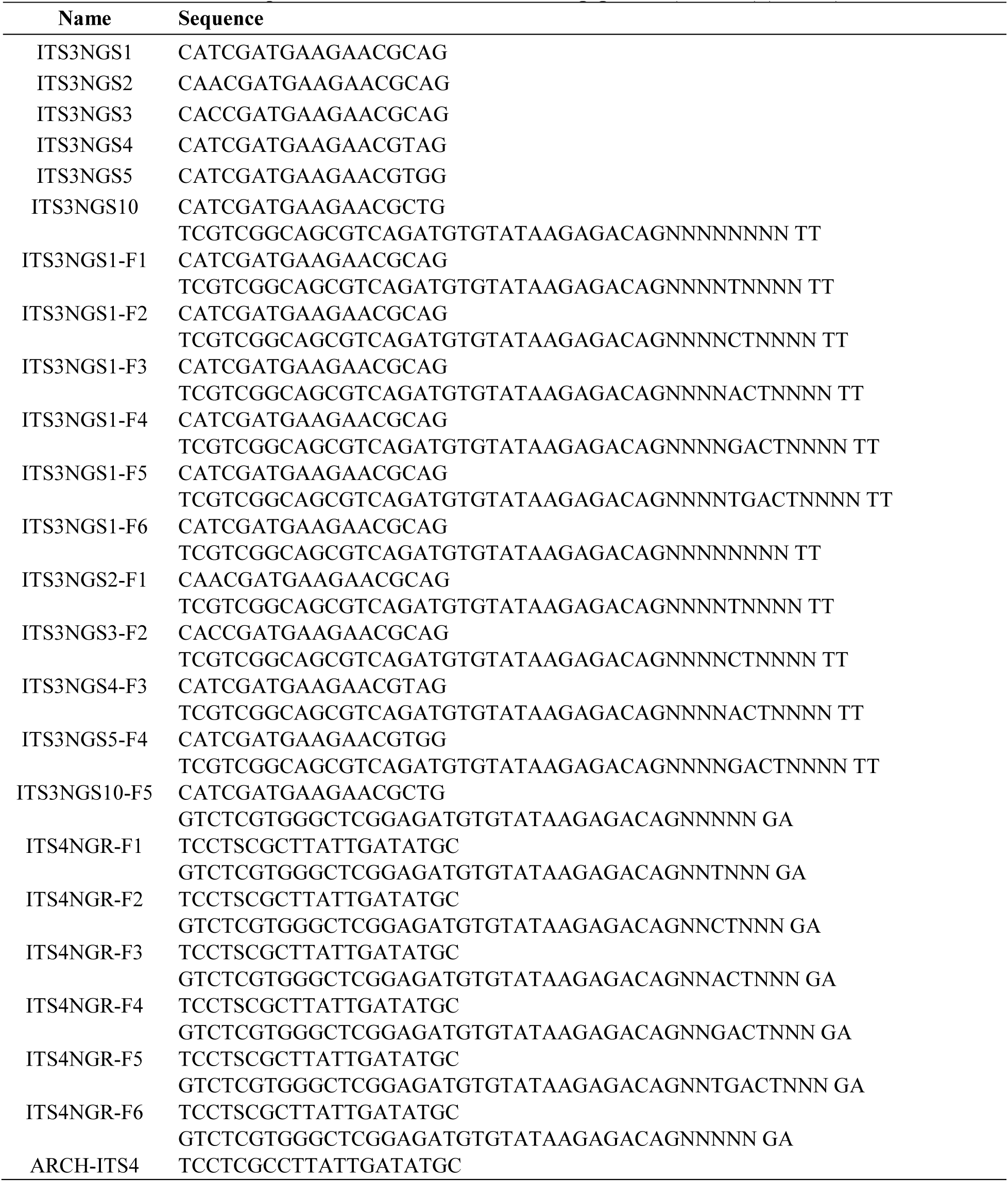
Primers and sequences used in Illumina ITS2 pipeline (5’ to 3’) (24–26).

**Table S3.** Heatmap (Fig. 3C) environmental health-related genera statistics.

| Detected Targeted Genera | Class | Wilcoxon test adjusted p-value |  |
| --- | --- | --- | --- |
|  |  | Overall | Subsets |
| Alternaria | Airborne allergen | 0.072 |  |
| Aspergillus | Opportunistic pathogen | 0.00003 | Air, 0.008<br>Swab, 0.04 |
| Aureobasidium | Opportunistic pathogen | 0.072 |  |
| Bipolaris | Plant pathogen | 0.087 |  |
| Bjerkandera | Wood-decay | 0.389 |  |
| Candida | Pathogen | 0.039 |  |
| Cladosporium | Allergen | 0.004 |  |
| Cryptococcus | Pathogen | 0.056 |  |
| Epicoccum | Indoor fungus | 0.917 |  |
| Fusarium | Plant pathogen | 0.0004 | Swab, 0.03 |
| Malassezia | Pathogen | 0.002 | Air, 0.04 |
| Mucor | Pathogen | 0.572 |  |
| Penicillium | Pathogen | 0.0008 | Air, 0.04 |
| Phoma | Plant pathogen | 0.057 |  |
| Rhodotorula | Opportunistic pathogen | 0.057 |  |
| Saccharomyces | Opportunistic pathogen | 0.015 |  |
| Stemphylium | Plant pathogen | 0.057 |  |
| Trichoderma | Opportunistic pathogen | 0.146 |  |
| Wallemia | Allergen | 0.052 |  |

\*Subsets are comparing between pipelines within select subsets of sample type (Air, Swab, Negative Control, Positive Control) or location (Home1, Home2, Lab 1, Lab 2, Dorm). Subsets with significant differences were listed.

## Supplementary Method

### Sample collection and preprocessing

#### Home 1 and Dorm

At Home 1 and Dorm, surface swabs were collected from a variety of surfaces using Isohelix™ SK-3S rayon swabs pre-moistened with 50 mM Tris buffer. Swabbing was performed over a 110 cm² area using standardized multidirectional S-strokes for 2 minutes. To account for potential contamination during field handling, field blanks were collected at both locations: two at Home 1 and one at Dorm. For each field blank, a sterile swab was opened in the field, briefly exposed to ambient air for 5 seconds, and placed directly into elution buffer without contacting any surface. Following sample and field blank collection, all swabs were transferred into an elution buffer (20 mM Tris, pH 8.0, 1% polyvinylpyrrolidone [PVP], 1% Tween 20) for transport. In the laboratory, swabs were briefly vortexed, and the resulting eluates were collected and aliquoted for downstream processing. All eluates were stored at −80 °C for no more than 6 months prior to analysis. At Home 1, bioaerosol samples (n = 8) were collected from three bedrooms using a BobCat (BC) AC-200 Sampler (InnovaPrep, Drexel, MO, USA), operated at 200 L/min for 2 hours with a dry electret filter. After sampling, the filter was eluted with 8 mL of wet foam (AC08100T, InnovaPrep). To control for potential contamination introduced during the aerosol collection and processing workflow, two method blanks were prepared by passing sterile wet foam through clean dry electret filters using the same procedure as for bioaerosol samples, but without field deployment. All eluates, including those from method blanks, were aliquoted and stored at −80 °C.

#### Home 2 and Lab

At Home 2 and Lab (1 and 2), surface swabs were collected using Isohelix™ SK-3S rayon swabs moistened with QIAGEN Buffer AE (10 mM Tris-Cl; 0.5 mM EDTA; pH 9.0). At Home 2, swabs were collected from a basement wall and an entryway wall. At UNC, swabs were taken from visibly mold-affected locations, including an air vent in a laboratory (Lab_1) and a light diffuser panel in a walk-in cold room (Lab_2). To monitor for potential contamination and assess processing consistency, one negative swab control (NC_swab) was included by exposing a sterile swab to ambient air in the laboratory for 5 seconds before placing it directly into QIAGEN Buffer AE without surface contact. One positive swab control (PC_swab) was prepared by spiking a sterile swab with Aspergillus niger. Following sample and control preparation, all swabs were transferred to QIAGEN Buffer AE for transport. In the laboratory, swabs were briefly vortexed, and the resulting eluates were collected and aliquoted for downstream processing. All eluates were stored at −80 °C for no more than 6 months prior to analysis. Bioaerosol samples were collected at both Home 2 and UNC using two complementary methods: (1) the BobCat (BC) as described above, operated at 200 L/min for 5 minutes; and (2) filter concentration, in which a 1 mL aliquot of the BC eluate was passed through a 47 mm, 0.45 µm mixed cellulose ester (MCE) membrane filter (Millipore HAWP 04700) to enhance particle retention. One negative bioaerosol control (NC_BM) was prepared by passing sterile wet foam through a clean dry electret filter without field deployment. One positive bioaerosol control (PC_BM) was prepared by spiking a sterile BobCat eluate with *Aspergillus niger* derived from ATCC 6275 (KwikStik). All eluates, including those from negative and positive controls, were aliquoted and stored at −80 °C until processing.

#### Illumina ITS2 library preparation PCR conditions

Fungal ITS2 regions were amplified using a three-step PCR protocol adapted from Lundberg et al. (23). Initial amplification of ITS2 was performed on 3µL of extracted total DNA in a 10-μL volume containing 0.3 μL of 10 μM forward (ITS3NGS) and reverse(ITS4NGR) primers (24–26), 5 μL of 2X KAPA HiFi HotStart ReadyMix (F. Hoffmann-La Roche AG, Basel, Switzerland), 0.1 μL of 100X SYBR Green I (Thermo Fisher Scientific, Waltham, MA, USA), and 1.3 μL nuclease-free water. Reaction mixes were denatured for 3 min at 95°C prior to 30 cycles of denaturation at 98°C for 20 seconds, annealing at 58°C for 15 seconds, and extension at 72°C for 1 min, followed by immediately cooling reactions to 4°C until transferred to ice. A second PCR introduced Illumina sequencing adaptors using frameshifted primers (ITS3NGS F1–F6, ITS4NGR F1–F6). 3µL of PCR product from first PCR was added to PCR master mix with same recipe as the previous PCR and ran at the same thermocycle conditions except for only repeat 10 cycles of denaturation-annealing-extension. Third PCR amplification to add Illumina adapters and dual 8-bp indices for sample multiplexing was performed in a 50 μl volume containing 5 μl of 2.5 μM forward and reverse indexing primers, 25 μl of 2X KAPA HiFi buffer, 0.5 μl of 100X SYBR Green I, 9.5 μl nuclease free water, and 5 μl of PCR product from previous step. Reaction mixes were denatured for 3 min at 95°C prior to 30 cycles of denaturation at 98°C for 20 seconds, annealing at 60°C for 15 seconds, and extension at 72°C for 1 min, followed by immediately cooling reactions to 4°C until transferred to ice. All primer sequences are listed in S. Table 2.

#### PacBio full-length ITS library preparation

Full-length ITS library was prepared using the same extracted DNA from ITS2 pipeline. Five µL of extracted total DNA was subject to PCR amplification of the ITS region using Phusion Plus PCR Master Mix (Thermo Fisher Scientific, Waltham, MA, USA) with forward primer ITS1catta (5’-ACCWGCGGARGGATCATTA-3’) and reverse primer ITS4ngsUni (5’-CCTSCSCTTANTDATATGC-3’) containing unique barcodes and Kinnex adaptors at a final concentration of 0.3uM(20, 27). Reaction mixes were denatured for 30 seconds at 98°C prior to 35 cycles of denaturation at 98°C for 10 seconds, annealing at 57°C for 20 seconds, and extension at 72°C for 75 seconds. After the last cycle, a final extension at 72°C for 5 minutes was performed. Completed PCR reactions were visualized on an Invitrogen E-Gel™ EX Agarose Gels (Thermo Fisher Scientific, Waltham, MA, USA) to ensure that amplicon size was correct (∼800bp), and that each sample amplified appropriately. Amplicon libraries were subsequently pooled based on gel band intensity (1.5, 5, 10, or 20µL per reaction). Library pool was cleaned and concentrated using 1.4x volume of SMRTbell® cleanup beads (Pacific Biosciences of California, Inc., Menlo Park, CA, USA) and eluted in 50µl of Low TE Elution Buffer (PacBio). Cleaned libraries were quantified using the Qubit HS dsDNA kit (Thermo Fisher Scientific, Waltham, MA, USA) and stored at -20°C prior to Kinnex PCR for concatenation and circularization. Kinnex PCR was performed as outlined in the PacBio Kinnex 16S kit’s published protocol with no modifications (PacBio)(28). Size selected and cleaned libraries were loaded onto a PacBio SMRT® Cell and sequenced on the Revio system (PacBio) in the Duke Sequencing and Genomic Technologies Shared Resource.

#### Illumina ITS2 Sequence Processing

Raw Illumina paired-end reads were processed using a multi-step pipeline combining several bioinformatic tools. First, paired-end reads were merged using PEAR (29) with default parameters to reconstruct full-length ITS2 amplicons. The ITS2 region was then extracted from merged reads using ITSxpress (30) with the following parameters: --region ITS2, --taxa Fungi, --single_end, and --threads 64. This step ensured that only the ITS2 region was retained for downstream analysis, removing flanking conserved regions.

Subsequent processing was performed using QIIME 2 (version 2023.9) (31) within a Singularity container environment. A manifest file was generated to import the ITSxpress-processed sequences into QIIME 2 as single-end data using the SingleEndFastqManifestPhred33V2 format. Quality control and amplicon sequence variant (ASV) inference were conducted using the DADA2 plugin with the denoise-single workflow (32). No length truncation was applied (--p-trunc-len 0) since sequences had already been quality processed through ITSxpress. The maximum expected error threshold was set to the default value of 2 (`--p-max-ee 2.0`). The DADA2 algorithm performed error correction, denoising, and chimera removal to generate high-resolution ASVs.

Taxonomic classification was performed using a pre-trained naive Bayes classifier based on the UNITE database (version 9.0, July 2023) obtained from the unite-train repository (33). The sklearn-based classifier was applied to representative ASV sequences using the classify-sklearn function with parallel processing enabled (--p-n-jobs -1). Final outputs including the feature table, representative sequences, and taxonomic assignments were exported from QIIME 2 artifacts using qiime tools export and converted to standard formats (TSV, FASTA) for downstream analysis in R.

#### PacBio full-length ITS Sequence Processing

PacBio circular consensus sequences (CCS) were processed using the DADA2 pipeline (version 1.28) with PacBio-specific modifications (32). Primer sequences (ITS1catta: 5’-ACCWGCGGARGGATCATTA-3’ and ITS4ngsUni: 5’-CCTSCSCTTANTDATATGC-3’) were removed using the removePrimers function with orientation correction enabled. Quality filtering and trimming were performed using filterAndTrim with the following parameters: minimum quality score of 3, minimum length of 250 bp, maximum length of 1200 bp, maximum expected errors of 3, and complete removal of sequences containing ambiguous nucleotides (maxN=0).

Error modeling was conducted using the PacBioErrfun function specifically designed for PacBio data, which accounts for the distinct error profile of long-read sequencing. Sequence variants were inferred using the dada function with PacBio-optimized error rates. Chimeric sequences were identified and removed using removeBimeraDenovo with the consensus method and a minimum fold-parent-over-abundance threshold of 3.5, which is optimized for longer amplicons.

Taxonomy was assigned using the same classifier based on the same UNITE database (version 9.0, July 2023) and sklearn-based classifier as Illumina short-read ITS2 sequences (33). The final amplicon sequence variant (ASV) table retained sequences ranging from 250-1200 bp, encompassing the full-length ITS region including ITS1, 5.8S rRNA gene, and ITS2.

#### Cross-platform ASV Mapping

The full-length ITS sequences and taxonomic assignments from the PacBio pipeline were extracted from the phyloseq object and exported to FASTA (pacbio_refseq.fasta) and tab-delimited taxonomy files (pacbio_taxonomy.tsv). To annotate Illumina-derived ITS2 amplicon sequence variants (ASVs), we used a direct sequence similarity search strategy. First, a nucleotide BLAST database was constructed from the PacBio reference sequences using makeblastdb from the NCBI BLAST+ suite. Illumina ITS2 representative sequences were then queried against this database using blastn, with a minimum sequence identity threshold of 97% and an e-value cutoff of 1e−20. The resulting alignment table was annotated with taxonomic information by joining the subject sequence identifiers (sseqid) with the PacBio taxonomy table using a custom R script. Final outputs included both the raw alignment metrics and the corresponding taxonomic assignments. We observed that several Illumina ASVs matched multiple full-length PacBio sequences, reflecting the limited resolving power of ITS2.

#### In-silico ITS2 Extraction from PacBio Full-Length ITS Reads

An in-silico ITS2 dataset was generated from the PacBio full-length ITS sequences (`pacbio_refseq.fasta`). The ITS2 region was extracted using `ITSx` with the --only ITS2 option to restrict output to the ITS2 subregion, and --taxa F to specify fungal ITS boundaries (34). The --preserve flag retained the original ASV identifiers, and the command generated the file `pacbio_ITSx.ITS2.fasta`. The resulting PacBio-ITS2 sequences were then processed and assigned with the same Illumina ITS2 pipeline to ensure classifier parity and identical QC/denoising settings. This PacBio-ITS2 dataset was used in sensitivity analyses to (i) compare rank-assignment rates to Illumina ITS2 under identical bioinformatic conditions, and (ii) quantify how much of the Full-length ITS vs ITS2 difference could be attributed to amplicon region and primer targeting rather than platform chemistry.

#### Statistical analysis

All statistical analyses and data visualizations were performed in R Studio Version 2024.04.2 (Posit, PBC, Boston, MA, USA). ITS2 sequencing depth sufficiency was evaluated by (i) rarefaction curve using function rarecurve and (ii) computing Good’s coverage per sample (1 − singletons/total reads). Before formal data analysis, functional guilds were added to fungal taxa in phyloseq objects using FUNGuild (49) with a confidence ranking of “Probable” or “Highly Probable”. Taxa were categorized into ecological guilds (saprotrophs, plant pathogens, animal pathogens, etc.) and trophic modes. To handle differences in sequencing depth, taxa abundance data underwent centered-log ratio (CLR) transformation using function transform from the microbiome package (35) to address the compositional nature of microbiome data. For visualization purposes, relative abundance transformations were also performed using the same package. Taxonomic data were agglomerated at various taxonomic ranks using the tax_glom function in phyloseq when appropriate.

Alpha diversity metrics including observed richness and Shannon diversity index were calculated using the vegan (37) package. Beta diversity was assessed using Bray-Curtis distances on relative abundance transformed data, ordination plot was using the Principal Coordinate Analysis (PCoA) in phyloseq package. Differences in fungal community composition between sample groups were tested using permutational multivariate analysis of variance (PERMANOVA) with the adonis2 function from vegan (37), using 10,000 permutations and Euclidean distances on CLR-transformed data. Ordination agreement was evaluated using symmetric Procrustes/PROTEST (vegan), reporting the Procrustes correlation 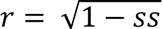 and permutation p-value, and Mantel (Spearman) on Bray–Curtis distance matrices. Analyses used matched samples and 999 permutations.

The taxonomic assignment success between Illumina and PacBio platforms was compared by calculating the percentage of ASVs assigned at each taxonomic level from kingdom through species. Correlations between platforms were assessed using Spearman’s rank correlation coefficients calculated on CLR-transformed abundance data at the genus level. Paired Wilcoxon tests -tests were performed to test differences between sequencing methods and between sample types in detecting environmental health related genera. Analysis of variance (ANOVA) was performed to test differences in per-sample correlation coefficients between sequencing platforms, followed by Tukey’s HSD post-hoc tests when significant differences were detected. Normality and homogeneity of variance assumptions were assessed using Shapiro-Wilk and Levene’s tests, respectively.

